# RNA-directed activation of cytoplasmic dynein-1 in reconstituted transport RNPs

**DOI:** 10.1101/273912

**Authors:** Mark A. McClintock, Carly I. Dix, Christopher M. Johnson, Stephen H. McLaughlin, Rory J. Maizels, Ha Thi Hoang, Simon L. Bullock

## Abstract

Polarised mRNA transport is a prevalent mechanism for spatial control of protein synthesis. However, the composition of transported ribonucleoprotein particles (RNPs) and the regulation of their movement are poorly understood. We have reconstituted microtubule minus end-directed transport of mRNAs using purified components. A Bicaudal-D (BicD) adaptor protein and the RNA-binding protein Egalitarian (Egl) are sufficient for long-distance mRNA transport by the dynein motor and its accessory complex dynactin, thus defining a minimal transport-competent RNP. Unexpectedly, the RNA is required for robust activation of dynein motility. We show that a *cis*-acting RNA localisation signal stabilises the interaction of Egl with BicD, which licenses the latter protein to recruit dynein and dynactin. Our data support a model for BicD activation based on RNA-induced occupancy of two Egl-binding sites on the BicD dimer. Scaffolding of adaptor protein assemblies by cargoes is an attractive mechanism for regulating intracellular transport.

## INTRODUCTION

Targeting of mRNAs to specific locations within the cytoplasm can confer precise spatial control over protein synthesis and function (Buxbaum et al, 2015; Holt & Schuman, 2013; Martin & Ephrussi, 2009). By compartmentalising protein function, mRNA localisation contributes to diverse processes, including embryonic axis determination, epithelial polarity and neuronal plasticity. Polarised trafficking of mRNAs frequently depends on the action of cytoskeletal motors, in particular those that translocate along the polarised microtubule network (Mofatteh & Bullock, 2017). However, the mechanisms by which specific mRNAs are recruited to, and transported by, microtubule motors remain unclear.

One of the most tractable systems for microtubule-based mRNA transport operates during early development of *Drosophila* melanogaster, and is responsible for localising spatial determinants of embryonic patterning to microtubule minus ends. Transport of these mRNAs is dependent on the Egalitarian (Egl) and Bicaudal-D (BicD) proteins (Bullock & Ish-Horowicz, 2001), as well as the minus end-directed motor cytoplasmic dynein-1 (dynein) and its accessory complex dynactin (Wilkie & Davis, 2001). Egl is a 1004-amino-acid protein that directly associates with the specialised RNA stem-loops that mediate polarised transport (so-called RNA localisation signals). The basis of RNA recognition is not known, although an exonuclease-like domain between residues 557 and 726 of Egl is partly responsible (Dienstbier et al, 2009). Egl uses a short N-terminal region to bind BicD (Dienstbier et al, 2009), and C-terminal features to bind the LC8 dynein light chain (Navarro et al, 2004). Mammalian BicD orthologues – BICD1 and BICD2 – associate with dynein and dynactin (Hoogenraad et al, 2001; Matanis et al, 2002). These observations have led to a model for linkage of localising mRNAs to the dynein transport machinery (Figure 1A). It is not known, however, if other factors co-operate with Egl and BicD to bridge mRNAs to the motor complex.

**Figure 1.**
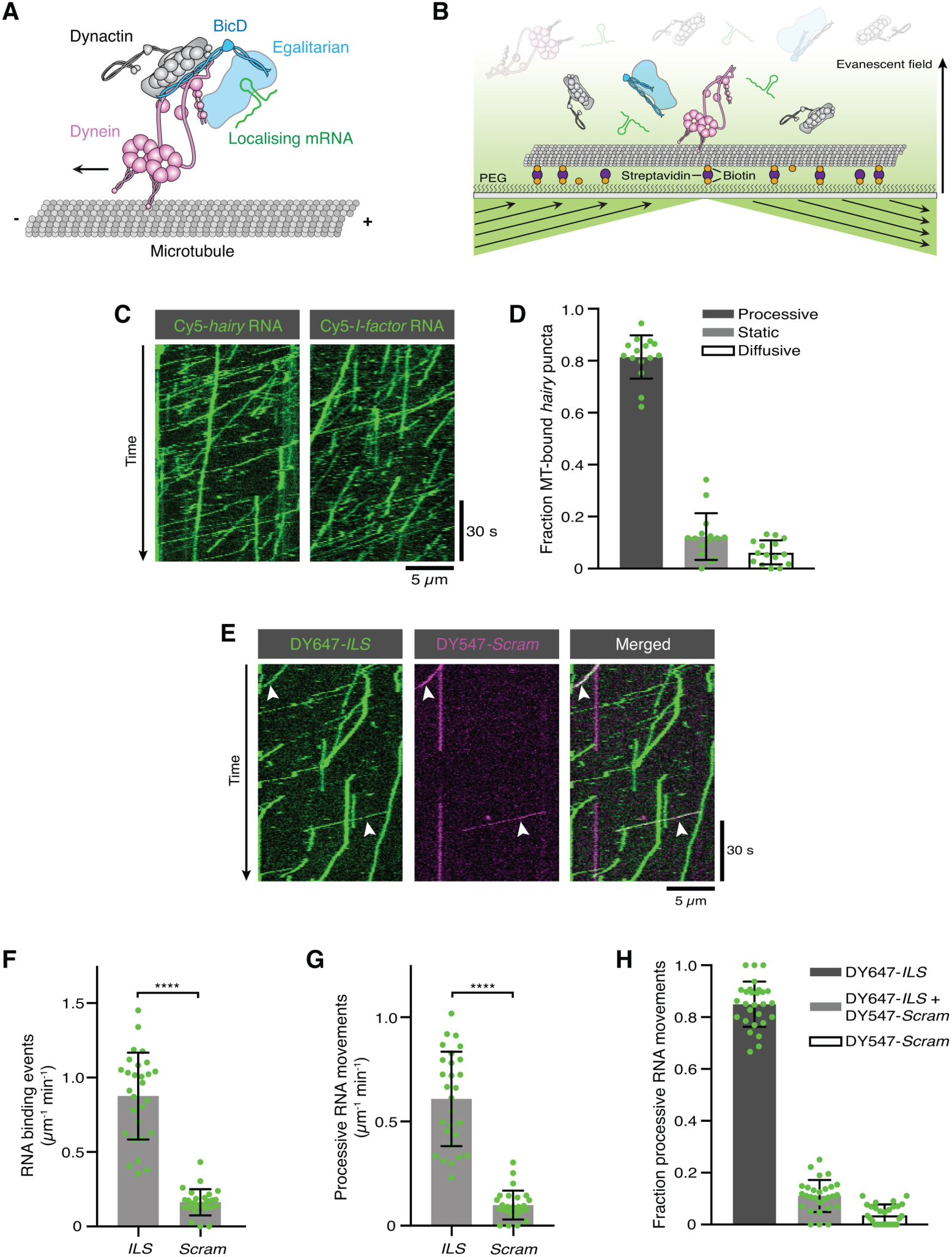
Reconstitution of dynein-based RNA transport with purified proteins. (A) Existing model for linkage of localising mRNAs to dynein. Note that there is no structural information available for Egl. (B) Diagram of TIRF-based *in vitro* motility assay. RNAs and proteins were mixed for at least one hour in the following molar concentrations: 100 nM dynein dimers, 200 nM dynactin, 100 nM Egl/BICD2 (with the operational assumption of two Egl molecules per BICD2 dimer) and 1 µM RNA. RNA-protein mixtures were typically diluted 40-fold and injected into imaging chambers containing microtubules that were pre-immobilised on passivated glass surfaces. (C) Examples of kymographs (time-distance plots) showing behaviour of Cy5-labelled *hairy* or *I-factor* RNAs in the presence of Egl/BICD2, dynein and dynactin. Diagonal lines are processive movements. In these and other kymographs, the microtubule minus end is to the left. (D) Fraction of microtubule (MT)-associated *hairy* RNA complexes that are processive, static or diffusive. (E) Kymograph illustrating behaviour of DY647-labelled *ILS* and a scrambled (*Scram*) version of the sequence labelled with DY547 following co-incubation with Egl/BICD2, dynein and dynactin. Arrowheads: examples of co-transport of the two RNA species. (F and G) Numbers of RNA binding events on microtubules (F) and processive RNA movements (G) of *ILS* and *Scram* RNAs. (H) Fraction of processive RNA movements that contain signals from the *ILS* only, *Scram* only, or both RNAs. In D and F-H, circles are values for individual microtubules. Error bars: SD. Statistical significance was evaluated with a Welch’s *t*-test (F and G). ****: P <0.0001.

Another outstanding question is how the assembly of the transport complex, and the activity of the dynein motor within it, is controlled. Several lines of evidence indicate that BicD is a key player in these processes. By forming an extended coiled-coil homodimer, the isolated N-terminal region of mammalian BICD2 (BICD2N: containing coiled-coil domain 1 (CC1) and part of CC2)) can bridge the interaction between dynein and dynactin, forming a mutually-dependent triple complex (Hoogenraad & Akhmanova, 2016; Splinter et al, 2012; Urnavicius et al, 2015; Zhang et al, 2017). The binding of dynein to BICD2N and dynactin increases the incidence of processive movement dramatically (McKenney et al, 2014; Schlager et al, 2014), which is associated with repositioning of the dynein motor domains with respect to the microtubule (Chowdhury et al, 2015; Zhang et al, 2017). The motor also moves with higher velocity and has increased force output once bound to BICD2N and dynactin (Belyy et al, 2016; McKenney et al, 2014). The equivalent N-terminal region of *Drosophila* BicD stimulates dynein-based transport *in vivo* (Dienstbier et al, 2009), suggesting that this mechanism is evolutionarily conserved.

Full-length BicD proteins interact poorly with dynein and dynactin and are therefore only weak activators of dynein motility (Dienstbier et al, 2009; Hoogenraad et al, 2001; Hoogenraad et al, 2003; Huynh & Vale, 2017; Liu et al, 2013). While mechanistic details are still lacking, BicD appears to be autoinhibited by folding back of the third coiled-coil domain (CC3) onto the dynein-activating sequences in CC1/2 (Dienstbier et al, 2009; Hoogenraad et al, 2001; Stuurman et al, 1999). It has been proposed that the interaction of CC3 with cargo-binding proteins such as Egl or Rab6 (a G-protein that binds Golgi-derived vesicles) frees CC1/2 to interact with dynein and dynactin (Dienstbier et al, 2009; Hoogenraad & Akhmanova, 2016; Hoogenraad et al, 2001; Hoogenraad et al, 2003; Matanis et al, 2002). Consistent with this model, mutating an essential residue in the shared Rab6-and Egl-binding site in CC3 prevents *Drosophila* BicD from associating with dynein *in vivo* (Liu et al, 2013).

It has recently been shown *in vitro* that the presence of Rab6 allows full-length BICD2 to associate with dynein and dynactin and thereby activate transport (Huynh & Vale, 2017). This observation provides direct evidence that association of a cargo-binding protein with CC3 stimulates the assembly of an active dynein-dynactin-BicD complex, although the stoichiometry of Rab6 and BICD2 in transport complexes was not investigated. Binding of mammalian and *Drosophila* BicD proteins to Rab6 is strictly dependent on the G-protein being GTP-bound (Rab6^GTP^) (Huynh & Vale, 2017; Liu et al, 2013; Matanis et al, 2002; Short et al, 2002), a state induced by association with its target membranes (Hutagalung & Novick, 2011). These data suggest a mechanism for linking long-distance movement of dynein with the availability of a vesicular cargo. Egl, on the other hand, can associate with BicD CC3 *in vitro* in the absence of an RNA cargo (Dienstbier et al, 2009; Liu et al, 2013). This observation implies that the RNA is not involved in the relief of BicD autoinhibition by Egl, although this hypothesis has not been tested directly.

We set out to elucidate molecular mechanisms of dynein-based mRNA transport by Egl and BicD by reconstituting this process *in vitro* with purified components. Our results define a minimal set of proteins for RNA translocation on microtubules and show that the RNA strongly activates motility of dynein. Stimulation of transport by RNA is not dependent on the Egl-LC8 interaction. Rather, our data support a model in which the RNA localisation signal overcomes BicD autoinhibition by stabilising the interaction of Egl with BicD CC3. Our study reveals a pivotal role of the RNA localisation signal in gating the activity of a microtubule motor, and give rise to a model in which cargoes activate dynein motility by scaffolding active adaptor protein assemblies.

## RESULTS

### An *in vitro* assay for dynein-based mRNA transport

We set out to determine if purified dynein, dynactin, Egl and BicD are sufficient to induce mRNA transport *in vitro*. As no method is available for the purification of *Drosophila* dynein and dynactin, we established a system in which *Drosophila* Egl and its mRNA targets are linked to mammalian dynein and dynactin complexes. We took advantage of the strong evolutionary conservation of the Egl/Rab6^GTP^-binding site of BicD (Figure S1; (Liu et al, 2013)) and produced a complex of *Drosophila* Egl bound to mouse BICD2. This complex was purified from Sf9 insect cells by co-expression of Egl with BICD2, as Egl was poorly expressed in the absence of its binding partner. The Egl/BICD2 complex, which was captured using an affinity tag on Egl, was not associated with significant amounts of RNA (Figure S2A). This observation is consistent with previous evidence that RNA is not essential for the interaction of Egl with *Drosophila* BicD (Dienstbier et al, 2009; Liu et al, 2013). The 1.4 MDa human dynein complex and 1.1 MDa pig dynactin complex were purified from established recombinant and native sources, respectively (Schlager et al, 2014). The purity of these and other protein preparations used in the study is illustrated in Figure S2B. RNAs were transcribed *in vitro*, and labelled by stochastic incorporation of fluorescent UTP.

Interactions of fluorescent RNA molecules with surface-immobilised microtubules were monitored by total internal reflection fluorescence (TIRF) microscopy in the presence of dynein, dynactin, and Egl/BICD2 (Figure 1B). RNAs and proteins were incubated together for at least one hour to promote complex assembly before dilution to sufficiently low concentrations to allow single molecules to be resolved on microtubules. We first used the 3’UTR of the *hairy* mRNA, which mediates transport by a complex containing Egl, BicD, dynein and dynactin during *Drosophila* embryogenesis (Bullock et al, 2003; Dix et al, 2013). We observed frequent association of *hairy* RNA with microtubules in the imaging chamber (Movie S1). Gratifyingly, 80% of microtubule-associated *hairy* mRNA puncta underwent long-distance transport (Figure 1C, D and Movie S1). As observed previously with a *Drosophila* extract-based system (Soundararajan & Bullock, 2014), *hairy* RNAs accumulated at microtubule minus ends following transport (Figure S3A) and were capable of diffusive motion on the microtubule lattice (Figure 1D and Figure S3B). We also performed experiments with the *I-factor* retrotransposon RNA, which is transported in association with Egl, BicD, dynein and dynactin during oogenesis (Dienstbier et al, 2009; Dix et al, 2013; Van De Bor et al, 2005). Like *hairy*, this RNA exhibited robust minus end-directed transport in our *in vitro* assay (Figure 1C). These experiments reveal that no additional proteins are required for microtubule-based mRNA transport *in vitro*. To test if RNA localisation signals are selectively recognised in our assay conditions we mixed the well-characterised 59-nucleotide Egl-binding element from the *I-factor* (*I-factor localisation signal* (*ILS*)) (Dienstbier et al, 2009; Van De Bor et al, 2005), which was labelled with DY647, with an equimolar amount of a scrambled version of the same sequence labelled with DY547.

In the presence of Egl, BICD2, dynein, and dynactin, the *ILS* bound to microtubules ∼five times more frequently than the mutant RNA and exhibited a similar relative increase in the number of processive movements (Figure 1E-G). Further analysis revealed that ∼75% of the processive complexes that contained the scrambled RNA also had a signal from the *ILS* (Figure 1E, H). This observation raises the possibility that much of the transport of the mutant RNA is due to association with the *ILS* rather than with Egl/BICD2. Collectively, these data reveal that the transport machinery retains selectivity for RNA localisation signals in our assay.

### Egl/BICD2 and dynactin are required for mRNA transport by dynein

We next investigated the involvement of each of the protein complexes in the RNA transport process. We first used SNAP tags to site-specifically label dynein and either Egl or BICD2 in the Egl/BICD2 complex. Egl and BICD2 were co-transported with dynein and the *hairy* RNA in the presence of dynactin (Figure 2A and Figure S4; note that dynactin could not be labelled as it is from a native source). Next, we omitted individual protein complexes from the assembly mix. The association of *hairy* with microtubules was barely detected when Egl/BICD2, dynactin, or dynein was excluded (Figure 2B, C and Figure S5). Thus, the simultaneous presence of all three complexes is required to link RNA to microtubules. In the absence of Egl/BICD2 or dynactin, dynein rarely exhibited transport but could still associate with microtubules (Figure 2B and Figure S5). However, there was an ∼two-fold increase in the frequency of microtubule binding events when both Egl/BICD2 and dynactin were present (Figure 2B, D). Thus, the combination of Egl/BICD2 and dynactin stimulates dynein’s ability to associate with microtubules and move processively in the presence of RNA.

**Figure 2.**
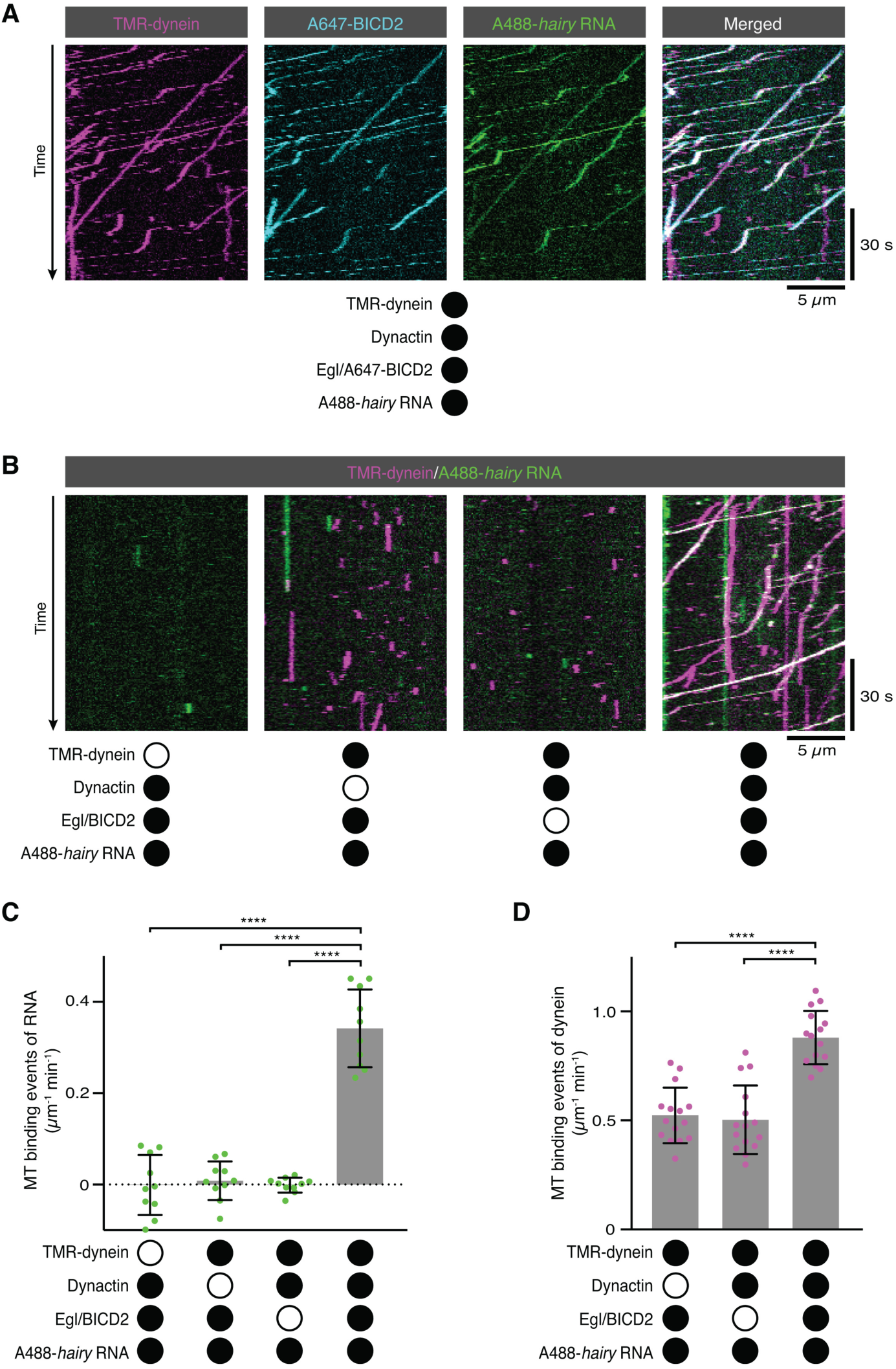
RNA transport by dynein requires the simultaneous presence of Egl/BICD2 and dynactin. (A) Kymographs showing co-transport of dynein, Egl (included in the assembly as a complex with unlabelled BICD2) and *hairy* mRNA in the presence of unlabelled dynactin. See Figure S4 for equivalent experiment with Egl labelled in the Egl/BICD2 complex. (B) Kymographs illustrating the results of omitting dynein, dynactin or Egl/BICD2 from the assay. Figure S5 shows images of separate channels. (C) Binding of Cy5-*hairy* RNA to microtubules in the presence of the indicated proteins. Signals were corrected for background binding of RNA to the glass surface. (D) Binding of TMR-dynein to microtubules in the presence of the indicated proteins. Background correction was not necessary due to negligible association of dynein with the glass surface. In this and other figures, black or white circles indicate proteins that were present or absent from the experiment, respectively. In C and D, small circles are values for individual microtubules. Error bars: SD. Statistical significance was evaluated with an ANOVA test with Dunnett’s multiple comparison correction. ****: P <0.0001

### RNA-directed activation of dynein motility

As described in the Introduction, the prevailing model is that the association of Egl with BicD CC3 is sufficient to free the N-terminal region of BicD to interact with dynein and dynactin. Unlike Rab6, Egl can bind BicD in the absence of associated cargo, leading us to ask whether dynein and dynactin differentiate between RNA-bound and RNA-free Egl/BICD2. To address this question, we performed motility assays with fluorescent dynein, dynactin and Egl/BICD2 in the presence and absence of RNA. Strikingly, the number of processive movements of dynein was ∼six-fold higher when the RNA was present (Figure 3A, B). This reflected an increase in both microtubule binding by dynein (Figure 3C) and the propensity for processive movement after engaging with the microtubule (Figure 3D). We also observed that the mean velocity and run length of dynein complexes bound to RNA were significantly higher than dynein complexes assayed in the absence of RNA (Figure 3E, F). We conclude that the RNA is required for robust stimulation of dynein motility and microtubule binding in the presence of Egl/BICD2 and dynactin.

**Figure 3.**
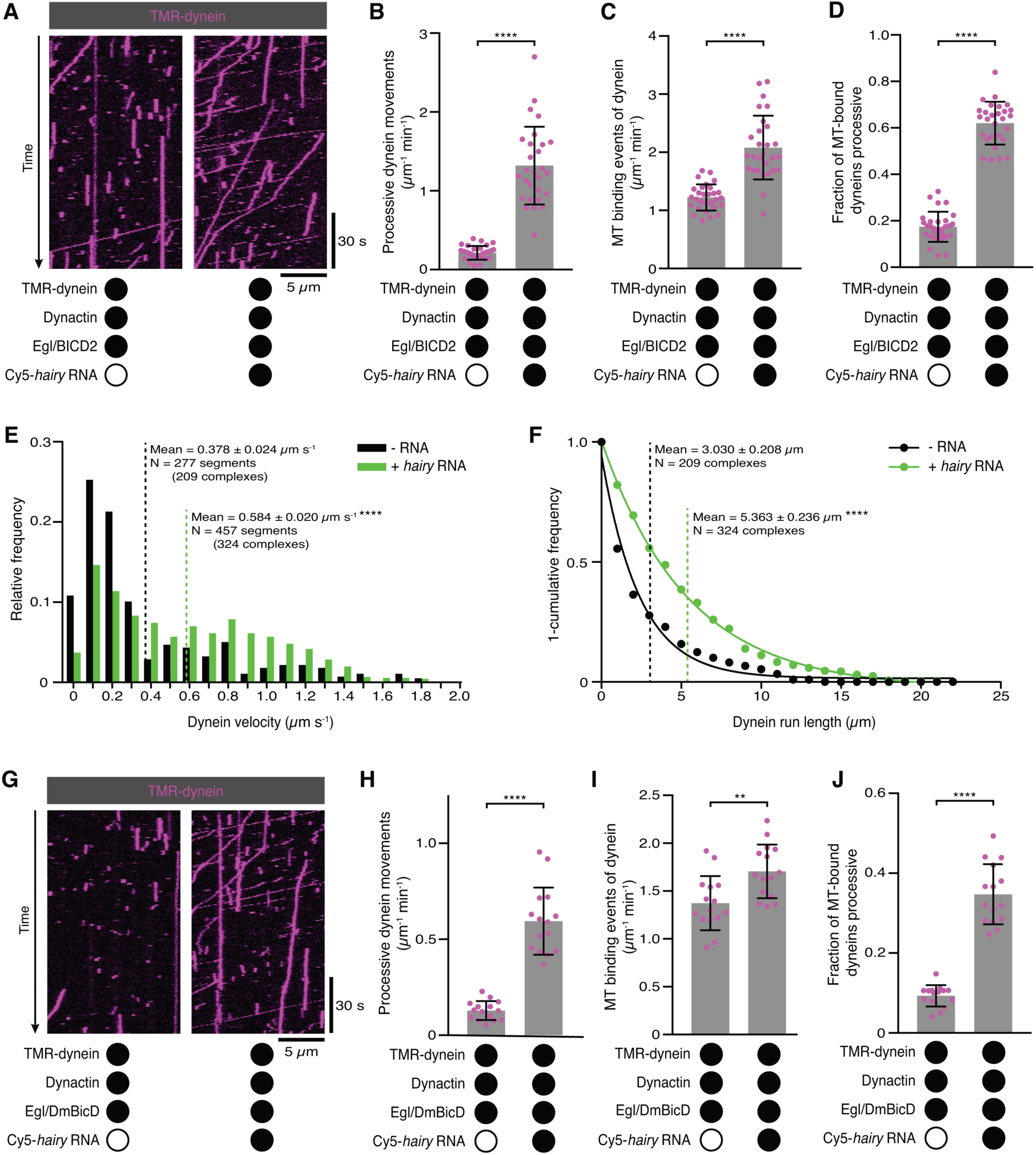
Activation of dynein motility by RNA. (A) Kymographs illustrating that *hairy* RNA increases the frequency of processive dynein movements in the presence of Egl/BICD2 and dynactin. (B-D) Numbers of processive dynein movements (B), dynein binding events on microtubules (C) and fraction of microtubule binding events that result in processive dynein movements (D) in the presence and absence of *hairy* RNA. (E and F) Distribution of segmental velocities (E) and run lengths (F) of dynein in the presence of Egl/BICD2 and dynactin ± *hairy* RNA (for experiments including *hairy*, only those complexes associated with the RNA were analysed). (G) Kymographs illustrating that *hairy* RNA increases the frequency of processive dynein movements when dynactin and a complex of Egl bound to *Drosophila* BicD (DmBicD) is included in the assay. (H-J) Numbers of processive dynein movements (H), dynein binding events on microtubules (I) and fraction of microtubule binding events that result in processive dynein movements (J) in the presence of dynactin and Egl/DmBicD ± *hairy* RNA. See Figure S6 for velocity and run length distributions for these experiments. Errors: SD, except in E and F (SEM). In B-D, and H-J, circles are values for individual microtubules. In B, C, H, and J, statistical significance was evaluated with a Welch’s *t-*test. In D and I, statistical significance was evaluated with a Student’s *t*-test. In E and F, statistical significance (compared to the equivalent parameter in the absence of RNA) was evaluated with a Mann-Whitney test using raw, unfitted values. **: P <0.01. ****: P <0.0001.

We next asked if the RNA-directed activation of dynein was a consequence of the combination of Egl with a BicD protein from a different species by performing experiments with a preparation of *Drosophila* Egl and *Drosophila* BicD (DmBicD). The Egl/DmBicD complex was also produced by co-expression of both proteins in Sf9 insect cells and purification with an affinity tag on Egl. The *hairy* RNA significantly increased the number of processive movements of dynein in the presence of dynactin when Egl/DmBicD was used (Figure 3G, H). This effect was again associated with enhanced microtubule binding of the motor, as well as increased probability of processive movement after engaging with the microtubule (Figure 3I, J). As was observed in the experiments with Egl/BICD2, the RNA also enhanced the mean velocity and length of dynein movements (Figure S6). Thus, RNA also gates the activation of dynein motility by a co-evolved Egl/BicD complex.

### RNA promotes the assembly of the Egl/BicD/dynein/dynactin complex

We considered two scenarios for how RNA stimulates dynein motility. First, the Egl/BicD/dynein/dynactin complex could be efficiently formed in the absence of RNA, with binding of RNA to Egl triggering a conformational change that activates processive dynein movement. Second, the ability of the Egl/BicD complex to interact with dynein and dynactin could be stimulated by the association of Egl with RNA, thus conferring different properties on the motor. To distinguish between these possibilities, we labelled Egl in the purified Egl/BICD2 complex with Tetramethylrhodamine (TMR) and monitored how *hairy* RNA affects its association with microtubule-bound Alexa647-dynein in the presence of dynactin. Although there was some association of dynein with Egl in the absence of RNA, the frequency of co-localisation increased by ∼six-fold when the RNA was present (Figure 4A-C). We confirmed that the RNA also stimulates the association of BICD2 with microtubule-associated dynein by labelling BICD2 within the purified Egl/BICD2 complex (Figure S7). Many dynein complexes that associated with Egl/BICD2 were motile (regardless of whether RNA was present of absent) (Figure 4B, Figure S7 and Figure S8), indicating that they were also bound to dynactin (McKenney et al, 2014; Schlager et al, 2014). Thus, the ability of RNA to activate processive dynein motion is associated with enhanced assembly of the Egl/BICD2/dynein/dynactin complex.

**Figure 4.**
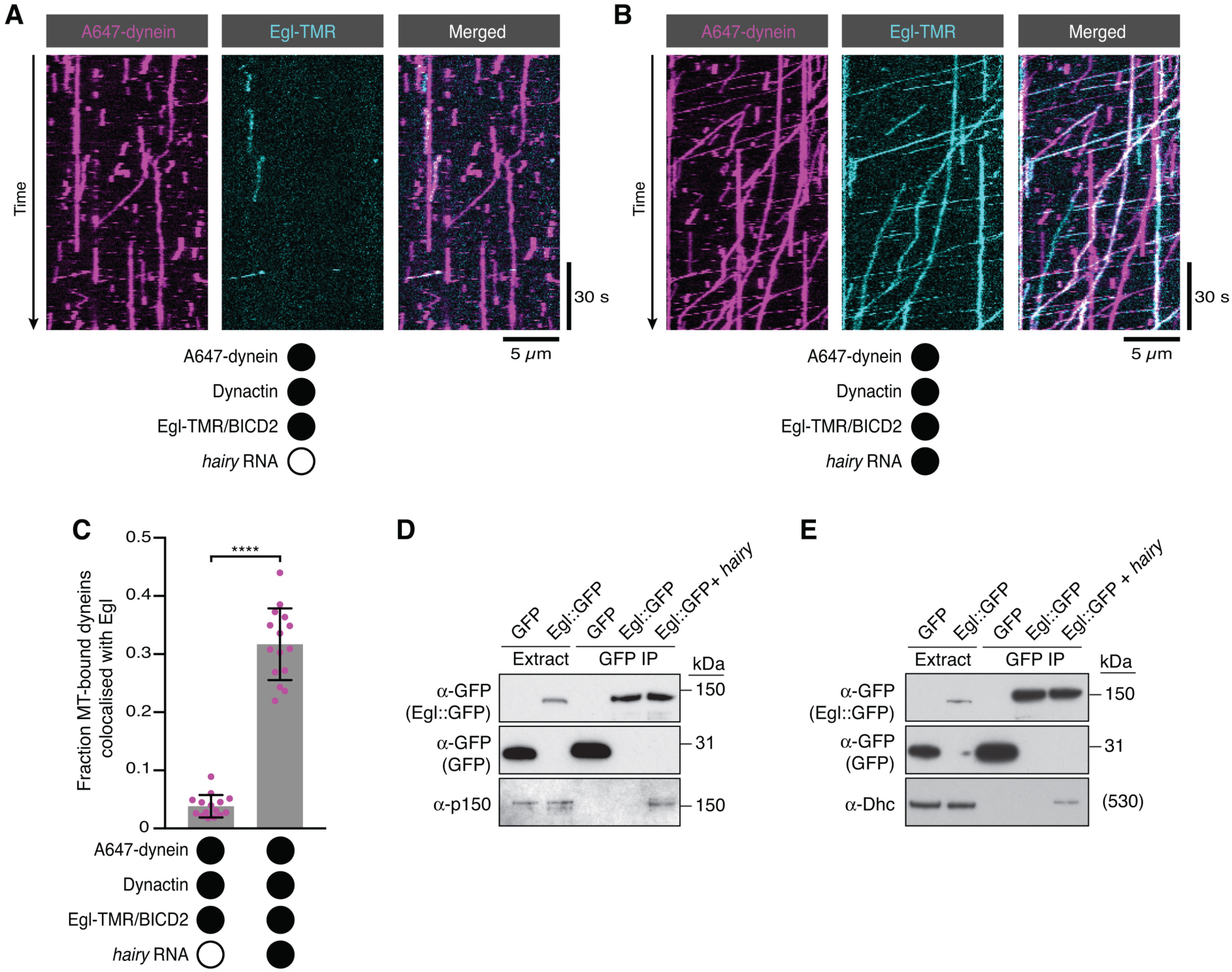
RNA stimulates the assembly of the transport complex. (A and B) Kymographs illustrating the behaviour of fluorescent dynein and Egl (included in the assembly in a complex with unlabelled BICD2) in the presence of dynactin ± *hairy* RNA. (C) Fraction of microtubule-associated dyneins that associate with Egl in the presence of dynactin ± *hairy* RNA. Circles are values for individual microtubules. Error bars: SD. Statistical significance was evaluated with a Welch’s *t-*test. ****: P <0.0001. See Figure S7 for equivalent data when BICD2 was labelled in the Egl/BICD2 complex. (D and E) Immunoblots of GFP-binding protein pulldowns from *Drosophila* embryo extracts showing RNA-induced co-precipitation of endogenous p150 (D) and Dhc (E) with Egl::GFP. This effect was observed in four independent experiments. For the blots shown, the amount of extract from which the loaded immunoprecipate was derived was 20 times the amount of extract loaded into the input lane for α-GFP, 200 times the amount of extract loaded into the input lane for α-Dhc and 1000 times the amount of extract loaded into the input lane for α-p150. Thus, only a small fraction of total Egl is associated with p150 and Dhc in the presence of RNA. Embryos expressing GFP were used as a control. In control experiments, the presence of RNA did not cause co-precipitation of the dynein-dynactin complex with GFP.

We next investigated if the assembly of the endogenous transport complex is stimulated by RNA. We immunoprecipitated a transgenically-expressed GFP-tagged Egl protein from *Drosophila* embryo extracts extracts in the presence and absence of exogenous *hairy* 3’UTR and assayed for co-precipitation of the p150 (DCTN1/Glued) subunit of dynactin and the heavy chain of dynein (Dhc) by western blotting (Figure 4D, E). p150 and Dhc were not detected in the Egl::GFP immunoprecipitate in the absence of exogenous RNA,. indicating that the association of Egl with dynein-dynactin is of low affinity or low abundance. In contrast, the addition of the *hairy* RNA led to detectable co-precipitation of the dynein and dynactin components with Egl::GFP. Thus, assembly of the transport complex is promoted by RNA in the context of both purified and endogenously-expressed proteins.

### The interaction of Egl with LC8 is not required for RNA-directed activation of dynein

The results described above raised the question of how RNA binding stimulates the association of Egl and the BicD protein with dynein and dynactin. We first asked if this involves the binding of Egl to LC8 dynein light chain. Motility assays were performed with a purified Egl/BICD2 complex in which Egl has two mutations in a consensus LC8-binding site that abolish association of LC8 *in vivo* and *in vitro* (Egl^dlc2pt^; S965K+S969R) (Navarro et al, 2004). The Egl^dlc2pt^/BICD2 complex supported robust transport of the *hairy* RNA in the presence of dynein and dynactin (Figure 5A and B). Moreover, in the presence of the mutant Egl/BICD2 complex and dynactin, the RNA still induced a strong increase in processive movement and microtubule binding of dynein (Figure 5C-E), as well as significantly higher mean velocities and run lengths of the motor (Figure S9). Thus, the interaction of Egl with LC8 does not play a significant role in the activation of transport by RNA.

**Figure 5.**
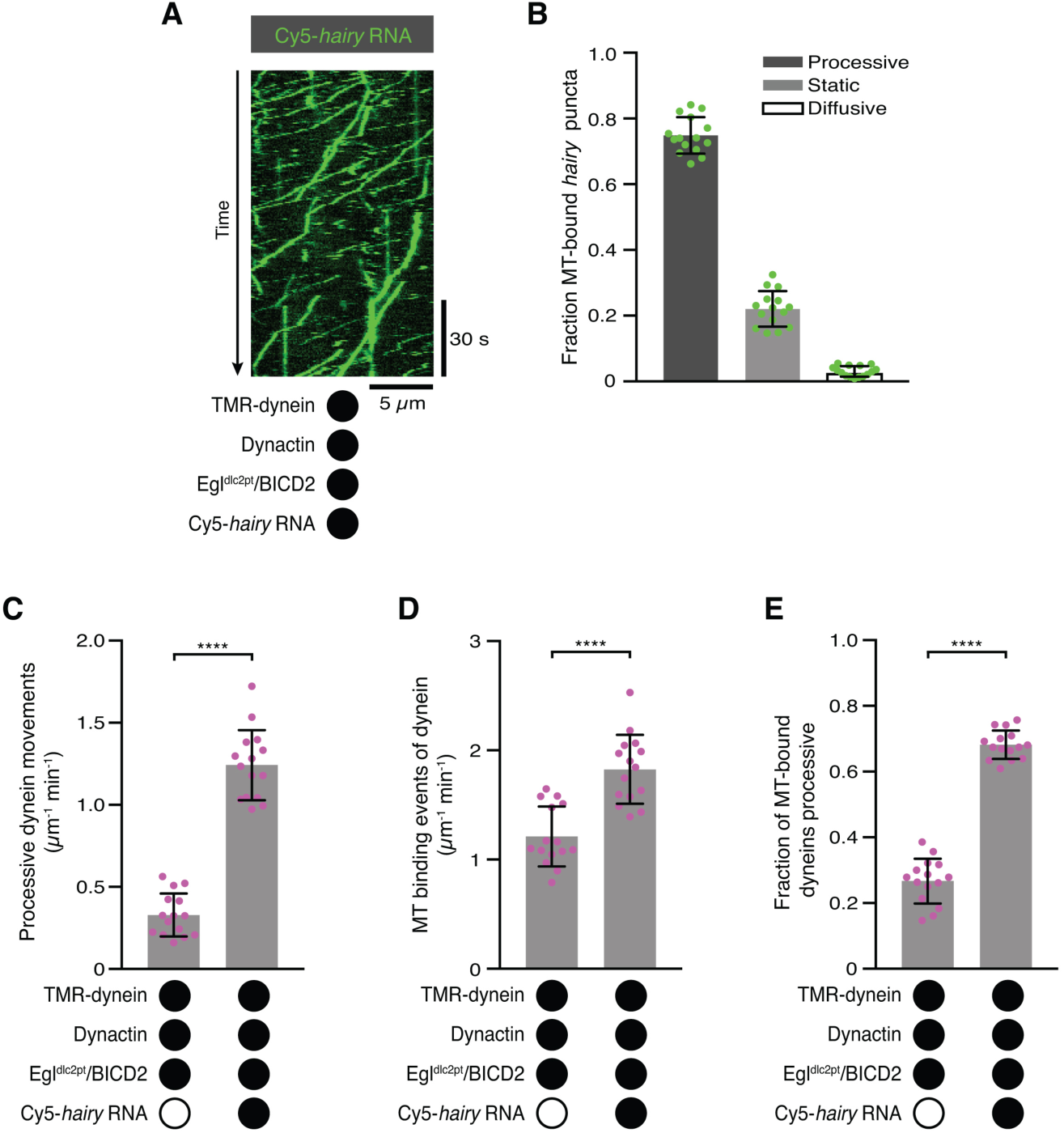
The Egl-LC8 interaction is dispensable for RNA-directed activation of dynein motility. (A) Kymograph illustrating robust transport of *hairy* RNA in the presence of dynein, dynactin and the Egl^dlc2pt^/BICD2 complex. (B) Fraction of microtubule-associated *hairy* RNA complexes that are processive, static or diffusive using the Egl^dlc2pt^/BICD2 complex. (C-E) Numbers of processive dynein movements (C), dynein binding events on microtubules (D) and fraction of microtubule-binding events that result in processive dynein movements (E) in the presence and absence of *hairy* RNA. In B-E, circles are values for individual microtubules. Error bars: SD. Statistical significance in C-E was evaluated with a Student’s *t*-test. ****: P<0.0001. See Figure S9 for velocity and run length distributions for these experiments.

### The RNA localisation signal stabilises the Egl/BicD complex

These observations pointed to the other reported interaction of Egl/BICD2 with dynein and dynactin, i.e. that mediated by BICD2N, as central to the activation of transport. As described in the Introduction, previous studies have indicated that occupancy of the Egl/Rab6^GTP^-binding site in BICD2 relieves autoinhibition, licensing BICD2N to interact with dynein and dynactin (Huynh & Vale, 2017; Liu et al, 2013).

During handling of the purified Egl/BICD2 complex, we noticed that it had a tendency to dissociate upon dilution. This observation suggests dynamic exchange of constituent species. We therefore wondered if the RNA relieves BICD2 autoinhibition by stabilising its interaction with Egl. To test this hypothesis, we mixed the 59-nt *ILS* RNA with purified Egl/BICD2 and performed size exclusion chromatography. The RNA localisation signal caused a large change in the elution profile of the protein complex compared to the RNA-free form (Figure S10), indicating a substantial increase in molar mass or a conformational change.

We next used sedimentation equilibrium analytical ultracentrifugation (SE-AUC) to evaluate mean molar masses of RNA-free and RNA-associated complexes independently of protein conformation. Over a range of protein concentrations, the presence of the *ILS* caused a large increase in mean molar mass compared to RNA-free samples (Figure 6A and Figure S11). An orthogonal method for determining molar masses – size-exclusion chromatography with multi-angle light scattering (SEC-MALS) – confirmed that the *ILS* increases the mean molar mass of the Egl/BICD2 sample substantially (Figure 6B and Figure S12). Gel-based analysis of the peak SEC-MALS fractions revealed that the *ILS*-induced mass increase was associated with more Egl in the fractions containing BICD2 (Figure 6B). We also used SEC-MALS to determine the effect of the *ILS* on the complex of Egl bound to DmBicD. The mean molar mass of the peak fractions increased substantially in the presence of the *ILS*, and this was again associated with increased association of Egl with the BicD protein (Figure 6C). Collectively, these experiments reveal that the Egl/BICD2 and Egl/DmBicD complexes readily equilibrate with constituent species and that this is counteracted by the presence of the RNA localisation signal.

**Figure 6.**
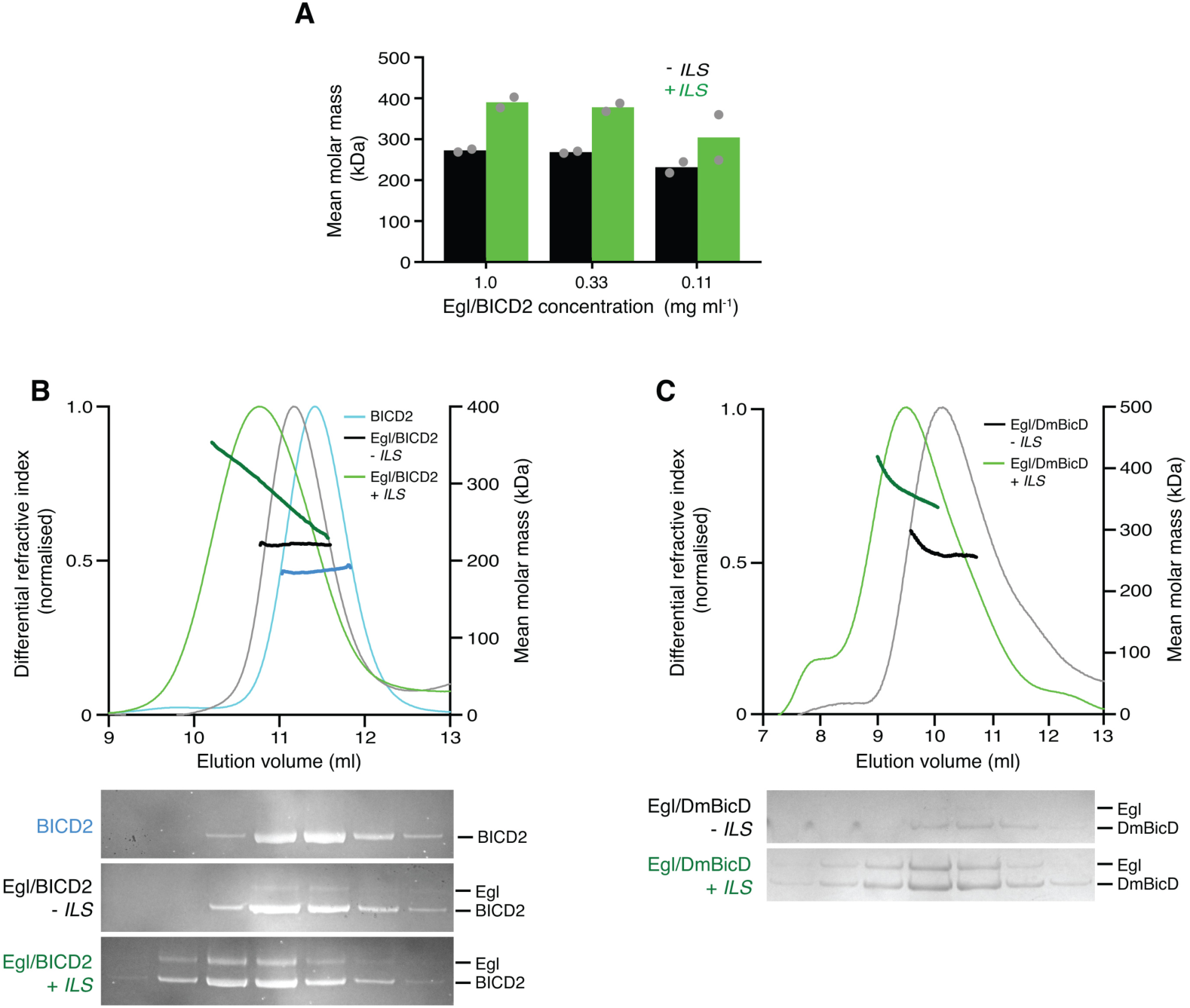
The RNA localisation signal promotes the occupancy of BicD with Egl. (A) Mean molar masses of Egl/BICD2 complexes in the presence and absence of the *ILS* RNA determined by SE-AUC. Circles are values for individual samples. See Figure S11 for examples of raw data and fitting. Protein concentrations of 1, 0.33 and 0.11 mg ml^-1^ equate to 24x, 8x and 2.7x the concentration in the assembly mix for *in vitro* motility assays. Experiments were performed at 4°C. (B and C) SEC-MALS analysis of sample of Egl/BICD2 (B) or Egl/DmBicD (C) in the presence and absence of *ILS* RNA. Darker lines indicate mean molar masses. The range of masses measured in the presence of the *ILS* indicates a mixture of multiple species. Free Egl elutes later from the column in a broad peak (not shown). Gels of collected fractions stained with SYPRO Ruby (B) or Coomassie (C) reveal more Egl associated with BICD2 or DmBicD in the presence of the *ILS*, which corresponds to species with higher molar mass (gels are aligned with corresponding positions in the SEC-MALS trace). Data for full length BICD2 alone are shown in B for comparison. SEC-MALS experiments with Egl/BICD2 were performed in 150 mM salt at room temperature, whereas those with Egl/DmBicD were performed in 50 mM salt at 4°C due to the relative instability of the complex. The concentration of the input samples was 0.5 mg ml^-1^. See Figure S12 for data for Egl/BICD2 using lower ionic strength buffer. In A-C, the *ILS* was always in a 10-fold molar excess to the protein (based on an assumption of two Egl molecules per dimer of BicD protein in the starting material).

The mean molar masses of Egl/BICD2 and Egl/DmBicD complexes determined by SEC-AUC and SEC-MALS ranged from ∼200 to 300 kDa in the absence of RNA depending on the experimental conditions, whereas the presence of the *ILS* led to mean molar masses of ∼300 to 450 kDa (Figure 6A-C; Figure S12). The predicted molar masses of the BICD2, DmBicD and Egl polypeptides are 93, 89 and 112 kDa, respectively, while the *ILS* has a molar mass of 19 kDa. It was previously shown that DmBicD is a dimer (Stuurman et al, 1999), and we confirmed that this is also the case for BICD2 using SEC-MALS (Figure 6B; observed molar mass 186.7 ± 0.5 kDa). The mean molar masses observed in our experiments are therefore compatible with the RNA promoting the occupancy of dimers of DmBicD or BICD2 with Egl, potentially culminating in a fraction of complexes containing two Egl molecules.

### The copy numbers of RNA, Egl and BicD in active transport complexes

To gain further insights into how RNA activates transport, we investigated the copy numbers of BICD2 and Egl in the complexes that are able to recruit dynein and dynactin and thus support processive movement on microtubules. We first produced Egl/BICD2 complexes that had SNAP tags on BICD2 and labelled them with a mixture of SNAP-reactive dyes such that approximately half of BICD2 polypeptides in the preparation were labelled with TMR, and approximately half were labelled with Alexa647. In an idealised situation, the exclusive presence of BICD2 dimers would result in 50% of complexes with one TMR dye and one Alexa647 dye, 25% with two TMR dyes and 25% with two Alexa647 dyes (Figure 7A).

**Figure 7.**
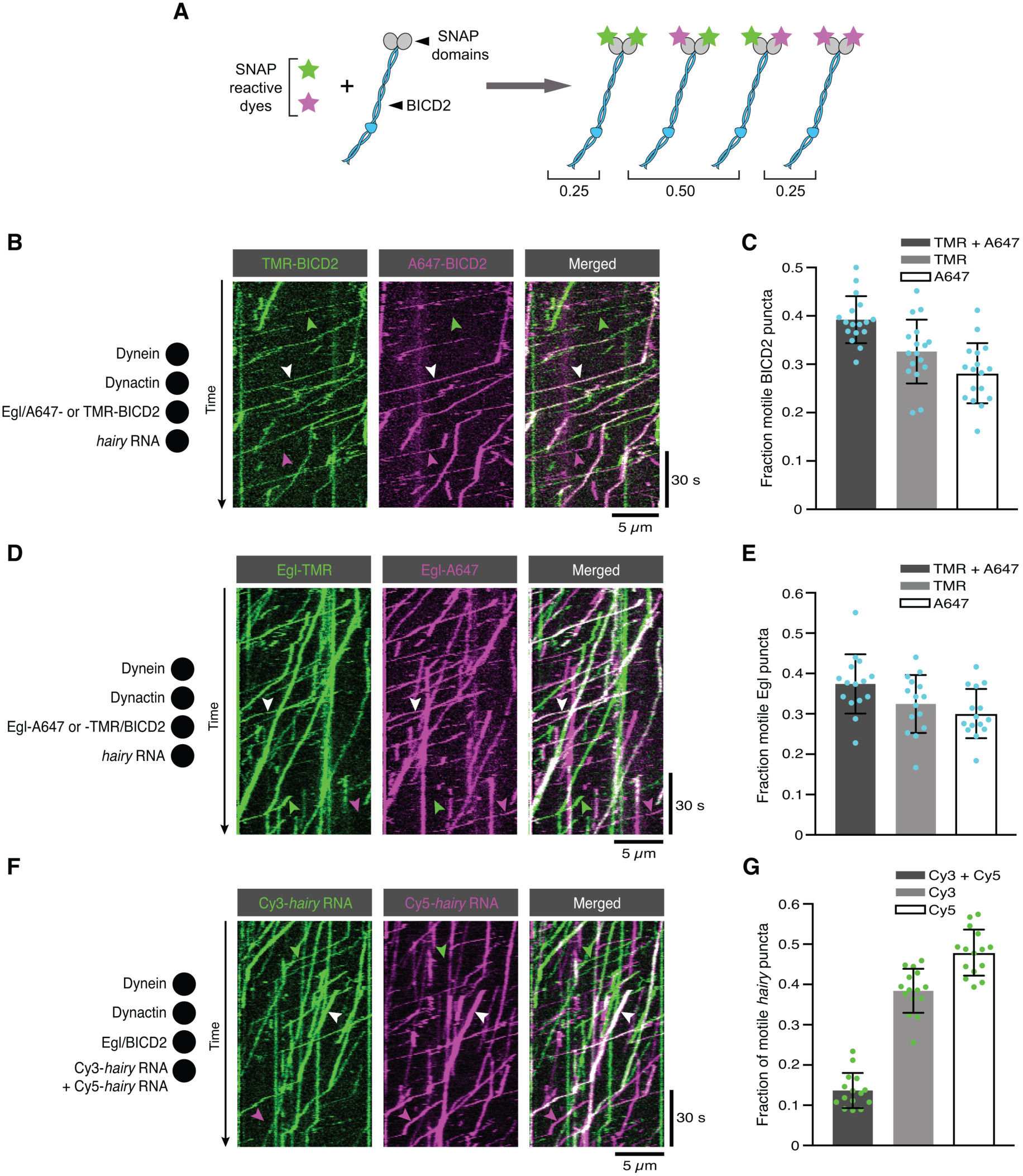
The copy number of BICD2, Egl and RNA in active transport complexes. (A) Idealised outcome of incubating a SNAP-tagged protein that is present in two copies per complex with equimolar amounts of two different SNAP-reactive dyes. The BICD2 dimer is used as an example, although the same principle applies for experiments with labelled Egl. (B) Kymograph of fluorescent signals when Egl/SNAP-BICD2 is labelled with a mixture of TMR and Alexa647 and assayed in the presence of RNA, dynactin and dynein. White, green and magenta arrowheads show, respectively, examples of motile BICD2-containing complexes with signals from both fluorophores, only TMR, or only Alexa647. (C) Fraction of motile BICD2-containing complexes with signals from both fluorophores, only TMR, or only Alexa647. (D) Kymograph of fluorescent signals when Egl-SNAP/BICD2 is labelled with a mixture of TMR and Alexa647 and assayed in the presence of RNA, dynactin and dynein. White, green and magenta arrowheads show, respectively, examples of motile Egl-containing complexes with signals from both fluorophores, only TMR, or only Alexa647. (E) Fraction of motile Egl-containing complexes labelled with signals from both fluorophores, only TMR, or only Alexa647. (F) Kymograph of fluorescent signals when Cy3-*hairy* and Cy5-*hairy* are mixed and assayed in the presence of Egl/BICD2, dynactin and dynein. White, green and magenta arrowheads show, respectively, examples of motile RNA-containing complexes with signals from both fluorophores, only Cy3, or only Cy5. (G) Fraction of motile *hairy* RNA puncta labelled with both fluorophores, only Cy3, or

When the labelled sample was used in motility assays with dynein, dynactin and *hairy* mRNA, 39% of the motile complexes with a BICD2 signal were labelled with both dyes (Figure 7B, C). Considering that the detection of dual coloured complexes will be preferentially affected by a small fraction of unlabelled polypeptides (see Methods), this result fits well with there being a single BICD2 dimer in most transport complexes.

When the procedure was repeated with SNAP-tagged Egl co-expressed with BICD2, the proportion of fluorescent complexes that was dual labelled in the presence of RNA was 37% (Figure 7D, E). This result indicates that there are two Egl molecules in the vast majority of active RNA transport complexes.

When this experiment was performed in the absence of RNA, the relatively small number of motile Egl complexes observed also had signal from both dyes in 41% of cases (Figure S13). These data indicate that even when the assembly of the transport complex is inefficient, motility is usually associated with the presence of two Egl molecules per complex. The capacity of BicD to bind two Egl molecules is compatible with the symmetrical nature of the Egl-binding region of CC3 (Liu et al, 2013).

Finally, we investigated the copy number of RNA in transport complexes by performing motility assays with or Cy5 dyes. We found that 14% of labelled motile RNPs had signal from both dyes (Figure 7F, G). Considering that a similar proportion of complexes will have two Cy3 dyes or two Cy5 dyes, these data suggest that at least 28% of RNPs contained two RNAs.

The observation that the frequency of dual-coloured RNAs is substantially lower than observed for BICD2 and Egl indicates that a sizeable fraction of transport complexes contain a single RNA molecule. We previously found that transport RNPs assembled in *Drosophila* extracts exclusively contain a single RNA (Amrute-Nayak & Bullock, 2012; Soundararajan & Bullock, 2014). The subset of complexes containing two *hairy* RNAs in our current assay may reflect non-specific RNA-RNA interactions that are normally blocked in extracts by the binding of other proteins.

In summary, the results of the dual-labelling experiments are consistent with the vast majority of BicD dimers in active transport complexes associating with two Egl monomers, and many associating with a single RNA. Together with the results of SE-AUC and SEC-MALS experiments, these data support a model in which the RNA localisation signal licenses BicD to bind dynein and only Cy5. In C, E and G, circles are values for individual microtubules; error bars: SD. dynactin by facilitating the association of two Egl molecules with BicD CC3 (see below).

## DISCUSSION

We have succeeded in reconstituting microtubule-based RNA transport *in vitro* using purified proteins and have used this system to define a minimal transport competent RNP. Although genetic experiments indicate that other proteins can modulate the mRNA transport process *in vivo* (Dix et al, 2013; Hain et al, 2014), no other factors appear to be obligatory for linkage of the RNA to dynein.

There has recently been considerable focus on the regulation of dynein motility, stemming from the discovery that the isolated N-terminal region of BicD proteins can bridge the interaction of dynein and dynactin and thereby activate transport (McKenney et al, 2014; Schlager et al, 2014; Splinter et al, 2012). Subsequent structural studies have provided important insights into how dynein activity is controlled in this system (Chowdhury et al, 2015; Grotjahn et al, 2018; Urnavicius et al, 2018; Urnavicius et al, 2015; Zhang et al, 2017). However, because full-length BicD proteins interact with dynein and dynactin poorly (Hoogenraad et al, 2001; Hoogenraad et al, 2003; Huynh & Vale, 2017), it is unclear how dynein activity is controlled within intact cargo-motor complexes.

Previous observations suggested that binding of a cargo-associated protein such as Egl to the C-terminal region of BicD is sufficient to overcome the autoinhibited state of the full-length protein, and thereby lead to recruitment of dynein and dynactin. Our study reveals that a purified Egl/BicD complex does not efficiently associate with dynein and dynactin in the absence of an RNA localisation signal. Thus, the availability of the cargo gates robust activation of dynein motility. This mechanism presumably limits unproductive long-range movement of the motor complex in the absence of an RNA consignment.

We show that the previously reported interaction of Egl with LC8 (Navarro et al, 2004) is not required for RNA-directed activation of dynein motility. Characterisation of the features of LC8 that mediate interaction with its binding partners also argue against a role for LC8 as an adaptor between Egl and dynein; the groove that LC8 uses to associate with the consensus binding motif present in Egl is also used for incorporation into the dynein complex, suggesting mutually exclusive interactions (Benison et al, 2007; Rapali et al, 2011). However, disrupting the Egl-LC8 interaction in *Drosophila* does significantly compromise the function of Egl in the maintenance of oocyte fate (Navarro et al, 2004). There are several examples of LC8 acting as a chaperone for binding partners, independently of its association with dynein (Rapali et al, 2011), and it may serve the same function for Egl *in vivo*.

Our data indicate that a key consequence of RNA binding to Egl is stimulation of the interaction of BicD with dynein and the processivity activator dynactin. Thus, RNA-bound Egl must overcome the autoinhibition of full-length BicD that prevents CC1/2 from interacting with dynein and dynactin. Negative stain electron microscopy in an accompanying preprint (Sladewski et al, 2018) lends further support to this notion; a folded back conformation of full-length DmBicD (Stuurman et al, 1999), which is likely to represent the autoinhibited state (Hoogenraad et al, 2001; Hoogenraad et al, 2003; Liu et al, 2013), was retained in the presence of Egl alone, but not detected in the presence of both RNA and Egl.

Our single molecule analysis is also consistent with activation of transport by RNA being mediated by CC1/2. The recruitment of RNA to dynein by Egl/BICD2 is dependent on dynactin (Figure 2B, C), as is also the case for the interaction of BICD2N with the motor complex (McKenney et al, 2014; Schlager et al. 2014; Splinter et al, 2012) Furthermore, the ability of RNA-bound Egl/BICD2 to augment dynein’s binding to microtubules (Figure 3C), as well as its velocity (Figure 3E) and run length (Figure 3F) is also shared with BICD2N (McKenney et al, 2014). Very recently, it has been shown that a single BICD2N and a single dynactin can recruit one or two dynein complexes (Grotjahn et al, 2018; Urnavicius et al, 2018), with the binding of the second motor increasing velocity and run length (Urnavicius et al, 2018). The two-motor state is associated with a subtle difference in the position of the N-terminal region of BICD2 CC1 (Urnavicius et al, 2018). Sladewski et al. (2018) provide evidence that two dynein motors are present in the majority of their transport RNPs, raising the possibility that interaction of RNA-bound Egl with BICD modulates the velocity and run length of transport complexes by favouring the two-motor-binding conformation of CC1.

An *in vitro* study of a yeast actin-based mRNA transport complex also reported stimulation of processive movement by the RNA cargo (Sladewski et al, 2013) (although an independent investigation of the same complex reported no influence of the RNA (Heym et al, 2013)). The mechanism that we and Sladewski et al. (2018) propose for RNA-mediated activation of dynein – involving relief of autoinhibition of an adaptor – is distinct from the one proposed for the yeast transport complex, which is based on RNA-dependent dimerisation of monomers of the myosin motor (Sladewski et al, 2013). Thus, multiple strategies may have evolved to co-ordinate the processivity of cytoskeletal motors with the availability of an RNA cargo.

How could binding of RNA to Egl relieve BicD autoinhibition? Although a complex of Egl bound to BICD2 can be purified in the absence of RNA following overexpression in cells, our data indicate that it readily dissociates into constituent species. The presence of the RNA localisation signal stabilises the interaction of Egl with BicD CC3, which is mediated by the first 79 amino acids of Egl and a 42-amino-acid region of BicD (Dienstbier et al, 2009; Liu et al, 2013). Although we cannot rule out additional mechanisms of RNA-mediated activation of BicD, the most parsimonious explanation for our data is that the RNA promotes the occupancy of CC3 with Egl, thereby freeing CC1/2 to interact with dynein and dynactin (Figure 8).

**Figure 8.**
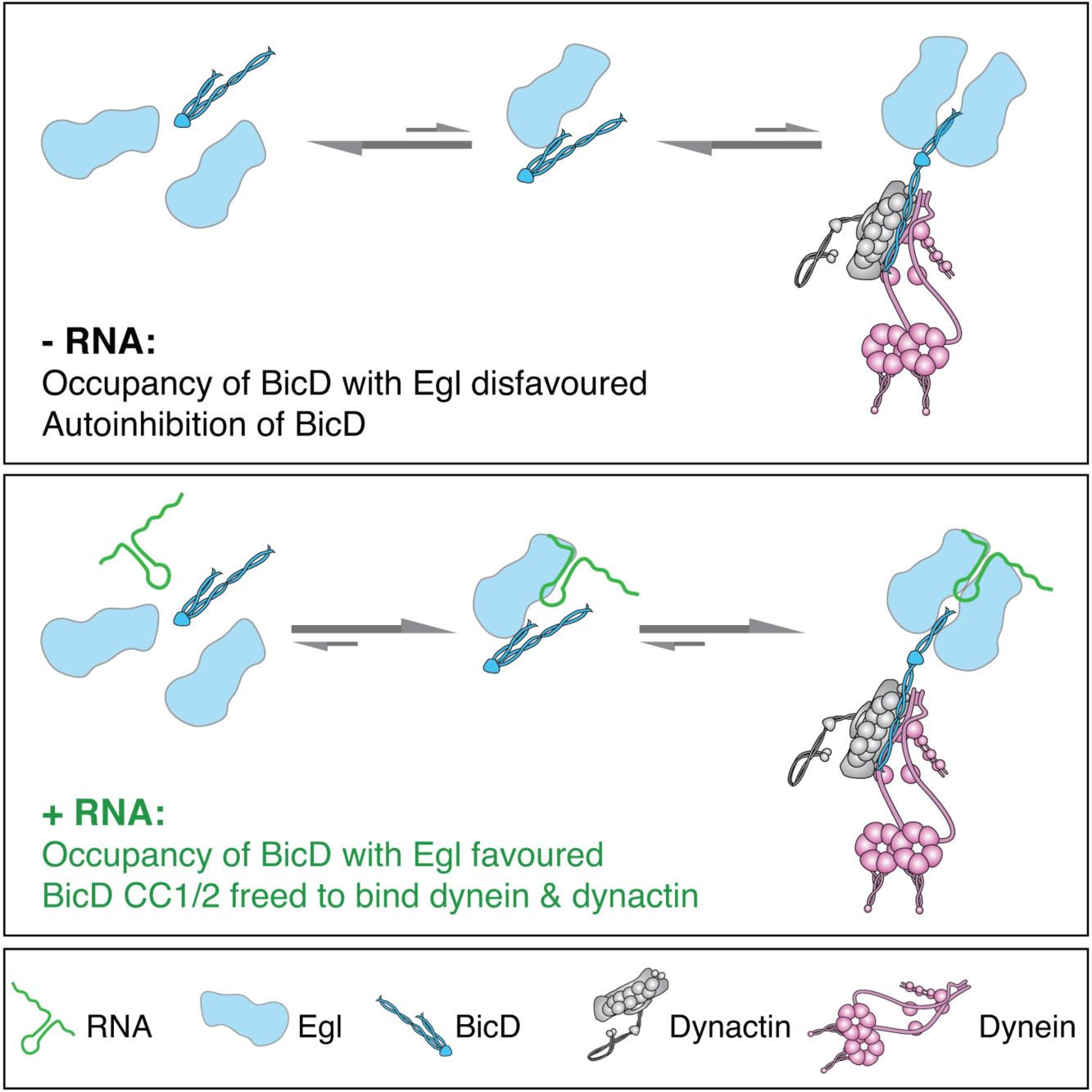
Model for the mechanism of RNA-stimulated assembly of an active dynein-dynactin complex. The RNA stabilises the interaction of Egl with CC3 of BicD, which promotes release of CC1/2 of BicD to interact with dynein and dynactin. We have depicted the structure of the Egl/BicD/dynein/dynactin complex equivalently in the presence and absence of RNA. It is, however, conceivable that in addition to stabilising the Egl/BicD complex, the RNA induces conformational changes in dynein and dynactin that also influence motility. A single RNA molecule is shown in the transport complex as our data indicate that this scenario is common.

It is not clear how binding of RNA-associated Egl (or Rab6^GTP^ for that matter) to BicD CC3 releases CC1/2. One possibility is that binding of Egl/Rab6^GTP^ competes directly with the interaction of CC3 with the N-terminal sequences. Alternatively, occupancy of the Rab6^GTP^-and Egl-binding site in CC3 could induce changes in coiled-coil architecture that are propagated along the molecule to release a discrete autoinhibitory interaction (Liu et al, 2013). The discovery of crystal forms of CC3 with different coiled-coil registers (Liu et al, 2013; Terawaki et al, 2015) lends support to the involvement of coiled-coil dynamics in the activation of BicD.

It is striking that the vast majority of active transport complexes contain two Egl polypeptides per BicD dimer, even when the transport process is compromised by the omission of RNA. This finding leads us to propose that occupancy of both Egl-binding sites of BicD CC3 favours the relief of BicD autoinhibition and recruitment of dynein and dynactin.

How could the RNA localisation signal stabilise the heterotetrameric Egl/BicD complex? Purified BicD was found not to interact directly with RNA localisation signals (Dienstbier et al, 2009), suggesting that the RNA does not act as bridge between Egl and BicD. Our single molecule experiments are consistent with two Egl proteins associating with a single RNA molecule. A structure-function study of an Egl-binding RNA localisation signal revealed two structurally-related helices that must be precisely registered with each other in order to trigger mRNA transport *in vivo* (Bullock et al, 2010). It is tempting to speculate that the two helices are discrete binding sites for Egl monomers, as this offers a simple explanation for how the RNA localisation signal facilitates the association of two Egl molecules with a BicD dimer. Alternatively, binding of the RNA localisation signal could induce a conformational change in Egl that increases the affinity for CC3, and thereby favours full occupancy of BicD. High-resolution structures of RNA-protein complexes will be required to discriminate between these possibilities.

In addition to Rab6^GTP^-associated vesicles and Egl-associated mRNAs, BicD proteins are implicated in transport of a diverse range of cellular cargoes and pathogens by dynein/dynactin (Dharan et al, 2017; Hoogenraad & Akhmanova, 2016; Indran et al, 2010; Redwine et al, 2017). It is conceivable that the cargo also promotes the activation of dynein in these systems by scaffolding the association of two cargo-associated proteins with CC3. It is easy to envisage how this could occur during membrane trafficking, when the diffusion of CC3-interacting proteins within the membrane would greatly facilitate self-association. The finding that BicD CC3 can simultaneously bind two Rab6^GTP^ proteins (Liu et al, 2013) is compatible with this scenario.

## METHODS

### Cloning and recombinant protein expression

Sequences encoding Egalitarian and BicD proteins (*Drosophila melanogaster* Egl isoform B:NM_166623, mouse BICD2:NM_001039179 and *Drosophila melanogaster* BicD:NM_165220) were synthesised commercially (Epoch Life Sciences, Sugar Land, TX) with codons optimised for expression in *Spodoptera frugiperda* Sf9 cells, and cloned for use with the MultiBac expression system. Where required, sequences encoding SNAPf tags for fluorescent labelling of protein complexes and ZZ-LTLT tags for IgG-based affinity purification (Reck-Peterson et al, 2006) were added by Gibson Assembly (NEB, Ipswich, MA) of PCR-amplified insert and backbone fragments. All constructs were validated by sequencing of the entire open reading frame. Genes encoding Egl-LTLT-ZZ or Egl-SNAPf-LTLT-ZZ were cloned downstream of the *polh* promoter of the pACEBac1 acceptor vector, while genes encoding BICD2, SNAPf-BICD2, or *Drosophila melanogaster* BicD (DmBicD) were cloned downstream of the *polh* promoter of the pIDC donor vector. The donor and acceptor vectors were recombined at defined Cre loci and subsequently incorporated into the baculovirus genome for simultaneous co-expression of Egl and BicD proteins. The same strategy was used for assembly of the gene encoding human DHC (tagged at the N-terminus with ZZ-LTLT-SNAPf) with those encoding other human dynein subunits, as described previously (Schlager et al, 2014). The isoform composition of the assembled dynein complex is as follows: DHC:NM_001376.4; DIC2:AF134477; DLIC2:NM_006141.2; Tctex:NM_006519.2; LC8:NM_003746.2 and Robl:NM_014183.3. All recombinant proteins were expressed from the baculovirus genome in Sf9 cells (Thermo Fisher Scientific, Waltham, MA) as described previously (Schlager et al, 2014). Cells were subsequently frozen in liquid N2 and stored at -80°C.

### Site-directed mutagenesis

The Egl^dlc2pt^ mutations (S965K + S969R) (Navarro et al, 2004) were generated by whole-vector PCR using a single pair of complementary mutagenic primers containing the desired sequence. Following amplification, the template DNA was digested with DpnI, and the amplicon ligated and propagated by transformation into α-Select Silver Efficiency chemically competent *E. coli* (Bioline, London, UK). The presence of the desired mutations, and no others, was confirmed by sequencing of the entire open reading frame.

### Protein purification

All purification steps were performed at 4°C. Native dynactin was purified from pig brain as described previously (Schlager et al, 2014; Urnavicius et al, 2015). Dynein, Egl/BICD2 and Egl/DmBicD complexes were affinity purified via an N-terminal ZZ-LTLT on DHC (ZZ-LTLT-SNAP_f_-DHC) and a C-terminal LTLT-ZZ tag on Egl (Egl-LTLT-ZZ or Egl-SNAPf-LTLT-ZZ). Frozen Sf9 cells were thawed on ice. For dynein purification, cells were resuspended in lysis buffer (50 mM HEPES pH 7.3, 100 mM NaCl, 10% glycerol, 1 mM DTT, 0.1 mM MgATP, 2 mM PMSF, 1 x cOmplete EDTA-free protease inhibitor cocktail (Sigma-Aldrich, St Louis, MO)). For purification of Egl/BICD2 and Egl/DmBicD complexes, lysis buffer was modified to include 500 mM NaCl to disrupt any association of Egl with native RNA species. Lysates were generated by repeated passage of resuspended cells through a Wheaton dounce tissue grinder (Fisher Scientific, Hampton, NH) and subsequently clarified by ultracentrifugation at 70,000 RPM (504,000 x *g*) using a Beckman Coulter Type 70 Ti fixed-angle rotor in a Beckman Coulter Optima L-100 XP preparative ultracentrifuge.

During centrifugation, IgG Sepharose 6 affinity resin (GE Healthcare Life Sciences, Little Chalfont, UK) was applied to a gravity flow Econo-column (Bio-Rad, Hercules, CA) and washed twice with 5 column volumes of lysis buffer (typical preps used 5 ml of resin slurry). Clarified lysate was added directly to the affinity matrix in the column, which was then sealed and agitated by gentle rolling for 3 h. After incubation, the lysate was allowed to flow through the column by gravity and the retained affinity matrix washed twice with 5 column volumes of lysis buffer and twice with 5 column volumes of TEV buffer (50 mM Tris-HCl pH 7.4, 150 mM KOAc, 2 mM MgOAc, 1 mM EGTA-KOH pH 7.3, 10% glycerol). If required, bound SNAPf-tagged proteins were fluorescently labelled on-column before proceeding to elution (see below). Bound proteins were eluted by overnight TEV cleavage of the ZZ affinity tag using gentle rolling agitation and ∼0.03 mg ml^-1^ TEV protease in a final volume of 15 ml TEV buffer. Eluted protein was recovered by gravity flow through a fresh Econo-column and concentrated to ∼1.5 mg ml^-1^ with a 100 kDa MWCO Amicon Ultra-4 centrifugal filter unit (Merck, Darmstadt, Germany).

The affinity-purified protein complexes were further purified by FPLC-based gel-filtration chromatography (AKTA Purifier and AKTA Micro, GE Healthcare Life Sciences) in GF150 buffer (25 mM HEPES pH 7.3, 150 mM KCl, 1mM MgCl_2_, 5 mM DTT, 0.1 mM MgATP, 10% glycerol) to remove large aggregates, TEV protease, and other small contaminants. For SEC-MALS and AUC experiments, GF150 was modified to include 5 mM TCEP instead of DTT. For the dynein complex, a TSKgel G4000SWxl with guard column (TOSOH Bioscience Ltd, Reading, UK) was used, while a Superose 6 Increase 3.2/300 column (GE Healthcare Life Sciences) was employed for Egl/BICD2 and Egl/DmBicD complexes. Fractions containing the dynein complex were pooled and concentrated to ∼1 mg ml^-1^. Fractions containing Egl/BICD2 or Egl/DmBicD complexes were pooled without an additional concentration step. All purified proteins were dispensed in aliquots for single-use, flash frozen in liquid N2, and stored at -80°C. Protein concentrations were determined using a Coomassie Protein Assay Kit (Thermo Fisher Scientific). To assess purity, proteins were resolved by SDS-PAGE using Novex 4-12% Bis-Tris precast gels (Thermo Fisher Scientific) and MES-SDS running buffer. Protein sizes were compared with Full-Range Rainbow prestained molecular weight markers (GE Healthcare Life Sciences). Protein bands were visualised using Coomassie Instant Blue protein stain (Expedeon, Over, UK) and imaged with a ChemiDoc XRS+ system (Bio-Rad).

### Fluorescent labelling of SNAPf-tagged proteins

Fluorescent labelling of SNAPf-tagged proteins with either SNAP-Cell TMR-Star (NEB) or SNAP-Surface Alexa Fluor 647 (NEB) was performed on-column during affinity capture according to a previously described method that labels >95% of dynein dimers with at least one dye (Schlager et al, 2014). For the mixed-labelling of SNAPf-BICD2 and Egl-SNAPf in Figure 7, an extended labelling time of 4 h and a further 10-fold excess of total SNAP-fluorophore reagent was used. This method labelled ∼ 90% of SNAPf-tagged polypeptides (∼80% of complexes containing two protein copies labelled with two dyes). Labelling efficiency was determined with spectrophotometry as previously described (Schlager et al, 2014). The ratio of SNAP-Surface Alexa Fluor 647 to SNAP-Cell TMR-Star that yielded approximately half of labelled polypeptides having one fluorophore and half the other fluorophore was determined empirically for different batches of the dyes.

### RNA synthesis and purification

Uncapped Cy5-*hairy* RNA or Cy3-*hairy* RNA was transcribed *in vitro* from a gel-purified PCR amplicon template using the MEGAscript T7 Transcription Kit (Ambion). The RNA is a 730-nt region of the 3’UTR containing the RNA localisation signal (Bullock et al, 2003). Cy5-UTP (PerkinElmer, Waltham, MA) was added to the transcription reaction together with a 4-fold excess of unlabelled UTP in order to label the RNA at multiple internal sites. Alexa488-*hairy* RNA was synthesised from the same template using a 1:9 ratio of Alexa488-UTP (Thermo Fisher Scientific) to unlabelled UTP. Cy5-*I-factor* RNA was synthesised from a linearised plasmid template using the MEGAscript SP6 Transcription Kit (Ambion) and a 1:3 ratio of Cy5-UTP to unlabelled UTP. The RNA is 597-nt long and contains the *ILS* localisation signal (Van De Bor et al, 2005). Following digestion of the template DNA with DNase, proteins were removed using phenol-chloroform-isoamyl alcohol (Thermo Fisher Scientific). Synthesised RNA was separated from unincorporated nucleotides by two rounds of purification with Sephadex G-50 size-exclusion spin columns (Sigma-Aldrich), precipitated with NH4OAc/ethanol and resuspended in nuclease-free dH_2_O. These procedures typically yield RNA samples with an average of ∼3 dyes per molecule. *ILS* wild-type (Van De Bor et al, 2005) and scrambled mutant RNAs (with and without a single 5’ DY547 or DY647 dye) were synthesised, decapped, deprotected, and HPLC purified by GE Dharmacon (Lafayette, CO). The mutant sequence was as follows (5’ to 3’):

AAAUGUGGUGCACUAUCUUCGUAUUCCAGUGCCACCGUGGUCUAAUUCACUCGUCGCC

An additional two A’s were included at the 5’ prime of synthetic RNAs to space the fluorophore from the wild-type or mutant localisation signal. The RNAs were further purified by gel-filtration chromatography in GF150 buffer (Superose 6 Increase 3.2/300, AKTA Micro (GE Healthcare)). All final RNA concentrations were determined using a Nanodrop 1000 spectrophotometer.

### Motility chamber preparation

Motility chambers with a volume of ∼10 µl were assembled by adhering glass cover slips functionalised with PEG/Biotin-PEG (Rapp Polymere, Tuebingen, Germany) to glass slides passivated with PLL-PEG (SuSos AG, Duebendorf, Switzerland) using three segments of double-sided tape distributed along the width of the slide. Glass surfaces were prepared as described previously (Bieling et al, 2010). The arrangement of tape yielded two parallel motility chambers per cover slip and allowed side-by-side comparison of two different conditions on the same glass surface. Prior to imaging, chamber surfaces were further passivated for 5 min with 1% (w/v) Pluronic F-127 (Sigma-Aldrich) and washed twice with 20 µl chilled motility buffer (30 mM HEPES pH 7.3, 5 mM MgSO_4_, 1 mM EGTA pH 7.3, 1 mM DTT, 0.5 mg ml-1 BSA). Chambers were then incubated with 2 mg ml^-1^ streptavidin for 5 min and again washed twice with 20 µl motility buffer. To block any unpassivated surface, chambers were incubated with 20 mg ml^-1^ α-casein (Sigma-Aldrich) for 5 min and washed twice with 20 µl motility buffer. The prepared chambers were kept in a humidified container until addition of microtubules and protein/RNA mixtures to prevent desiccation of chamber surfaces.

### Polymerisation and stabilisation of microtubules

Microtubules were polymerised from commercial porcine tubulin (Cytoskeleton Inc., Denver, CO) and labelled with fluorophores and biotin by stochastic incorporation of labelled dimers into the microtubule lattice. Mixes of 1.66 µM unlabelled tubulin, 0.15 µM Hilyte488-tubulin, and 0.4 µM biotin-tubulin were incubated in BRB80 (80 mM PIPES pH 6.85, 2 mM MgCl_2_, 0.5 mM EGTA, 1 mM DTT) with 0.5 mM GMPCPP (Jena Bioscience, Jena, Germany) for 2-4 h at 37°C. Polymerised microtubules were pelleted in a room temperature table top centrifuge at 18,400 x *g* for 8.5 min, and washed once with pre-warmed (37°C) BRB80. After pelleting once more, the microtubules were gently resuspended in pre-warmed (37°C) BRB80 containing 40 µM paclitaxel (taxol; Sigma-Aldrich) and used on the same day.

### In vitro motility assay

Constituents of motility assays were incubated together on ice for 1-2 hours by dilution into motility buffer to the following concentrations: 100 nM dynein, 200 nM dynactin, 100 nM Egl/BICD2 or Egl/DmBicD (using the operational assumption of two Egl molecules and one dimer of the BicD protein per complex), and 1 µM RNA. To ensure that all complexes assemble at the same ionic strengths, KCl was supplemented to a final concentration of 50 mM during assembly. Just prior to imaging, stabilised microtubules were immobilised in a prepared motility chamber for 5 min and subsequently washed once with motility buffer containing 50 mM KCl, 1 mg ml^-1^ α-casein, and 20 µM taxol. Assembly mixes were then diluted 40-fold (with the exception of the complexes in Figure 4A-C, which were diluted 20-fold) in motility buffer that additionally contained 50 mM KCl, 1 mg ml^-1^ α-casein, 20 µM taxol, 2.5 mM MgATP, and an oxygen scavenging system (1.25 µM glucose oxidase, 140 nM catalase, 71 mM 2-mercaptoethanol, 25mM glucose). This dilution was applied to the immobilised microtubules in the motility chamber for imaging at room temperature (23 ± 1°C).

### TIRF microscopy

For each chamber, a single multicolour acquisition of 500 frames was acquired at the maximum achievable frame rate (∼2 frames s^-1^) using a Nikon TIRF microscope system controlled with Micro-Manager open-source acquisition software (Edelstein et al, 2010) and equipped with a Nikon 100× oil objective (APO TIRF, 1.49 NA oil). The following lasers were used: Coherent Sapphire 488 nm (150 mW), Coherent Sapphire 561 nm (150 mW), Coherent CUBE 641 nm (100 mW). Images were captured with an iXon^EM^+ DU-897E EMCCD camera (Andor, Belfast, UK), resulting in pixel dimensions of 105 nm x 105 nm. Multicolour acquisitions used sequential image capture with rapid switching of emission filters (GFP, Cy3, and Cy5 (Chroma Technology Corp., Bellows Falls, VT)).

### Immunoprecipitation from Drosophila extracts

Extracts were generated from embryos of *P[tub-Egl::GFP]* (Dienstbier et al, 2009) or *Sco/CyO P[actin5C-GFP]* flies (Reichhart & Ferrandon, 1998), which contain genomically-integrated transgenes expressing Egl::GFP or GFP from the ubiquitous *α-tubulin* or *β-actin* promoters, respectively. 0-12 h embryos were dechorionated and flash frozen in liquid N_2_. 300 µl chilled extraction buffer (25 mM HEPES pH 7.3, 50 mM KCl, 1mM MgCl_2_, 2mM DTT, 2x cOmplete EDTA-free protease inhibitor) was added for each 100 mg of frozen embryos, followed by grinding on ice with a motorised pellet pestle (Fisher Scientific). The material was subjected to 25 passes in a Wheaton dounce tissue grinder (Fisher Scientific) on ice before the addition of 200 µl chilled extraction buffer containing 0.5% Triton-X-100 per 100 mg of embryos. Following gentle mixing, samples were incubated on ice for 5 min and passed through a 23G syringe five times before clarification by two centrifugation steps (each 5 min at 3000 x *g*). 350 µl aliquots of clarified extract were incubated with 20 units Recombinant RNAse Inhibitor (Promega, Madison, WI) and either 20 µl of a 6.7 µg/µl solution of unlabelled *hairy* RNA in dH_2_O or 20 µl dH_2_O for 30 min at 4°C. Magnetic beads coupled to GFP-binding protein (GFP-Trap MA (Chromotek, Martinsried, Germany)) were washed twice in PBS, followed by blocking of non-specific interaction sites with 1 mg ml^-1^ casein in PBS for 30 min at 4°C. After two washes of the beads in extraction buffer, the equivalent of 30 µl of initial bead slurry was mixed with the embryo extracts with or without *hairy* RNA. Following a 2 h 30 min incubation at 4°C, beads were washed fives times for 1 min in extraction buffer containing 0.05% Triton-X-100 (three washes in 400 µl of buffer and two washes in 1 ml buffer). Proteins and RNA-protein complexes were eluted from the beads by the addition of 60 µl 1 x lithium dodecyl sulphate (LDS) buffer (Thermo Fisher Scientific)/50 mM DTT and incubation at 80°C for 10 min.

Following electrophoresis and blotting onto PVDF membranes, proteins were detected using the following primary antibodies: mouse α-GFP (mix of clones 7.1 and 13.1 (Sigma-Aldrich); diluted 1:1000); mouse α-Dhc (clone 2C11-C (Sharp et al, 2000)) (both provided by the Developmental Studies Hybridoma Bank (University of Iowa, Iowa, IA) and diluted 1:1000) and rabbit α-p150-C-term ((Kim et al, 2007); provided by V. Gelfand, Northwestern University; diluted 1:10,000). Secondary antibodies were conjugated to horseradish peroxidase, with signal detected using the ECL Prime system (GE Healthcare) and Super RX-N medical X-ray film (FUJIFILM, Bedford, UK).

### Analytical ultracentrifugation

Duplicate independent preparations of 1 mg ml^-1^ Egl/BICD2 (2.4 µM assuming two Egl molecules and a single BICD2 dimer per complex) in GF150 buffer (using 5 mM TCEP instead of 5 mM DTT) in the presence or absence of a 10-fold molar excess of *ILS* RNA were pre-incubated on ice for at least 1 h and subsequently diluted in GF150 (TCEP) to yield 3 samples with volumes of 110 µl and protein concentrations of 1 mg ml^-1^, 0.33 mg ml^-1^, and 0.11 mg ml^-1^. These samples were loaded in 12 mm 6-sector cells and subjected to equilibrium sedimentation in an An50Ti rotor using an Optima XL-I analytical ultracentrifuge (Beckmann) at 3200, 5600, and 10,000 rpm until equilibrium was reached at 4°C. At each speed, comparison of several scans was used to judge whether equilibrium had been reached. Data were processed and analysed using SEDPHAT 13b (Schuck, 2003) and plotted with GUSSI (Brautigam, 2015). The partial-specific volumes (v-bar), solvent density and viscosity were calculated using Sednterp (T. Laue, University of New Hampshire).

### SEC-MALS

Samples of Egl/BICD2 and Egl/DmBicD at a starting concentration of 0.5 mg ml^-1^ were resolved on a Superdex 200 HR10/300 analytical gel filtration column (GE Healthcare) at 0.5 ml min^-1^ in GF150 buffer (using 5 mM TCEP instead of 5 mM DTT), GF75 buffer (contains 75 mM KCl), or GF50 buffer (contains 50 mM KCl). All measurements for Egl/BICD2 were made at room temperature, whereas the relative instability of the Egl/DmBicD complex required measurements be made at 4°C. Where indicated, *ILS* RNA was added at a 10-fold molar excess over Egl/BICD2 or Egl/DmBicD (based on two Egl molecules and a dimer of the BicD protein per complex) and incubated on ice for 1 h prior to injection on the column. Samples lacking RNA were treated in the same way. Following SEC fractionation, eluted protein was detected on a Wyatt Heleos II 18 angle light scattering instrument coupled to a Wyatt Optilab rEX online refractive index detector in a standard SEC-MALS format. Heleos detector 12 at 99° was replaced with Wyatt’s QELS detector for on-line dynamic light scattering measurements. Protein concentration was determined from the excess differential refractive index based on 0.186 RI increment for 1 g ml^-1^ protein solution. Concentrations and observed scattered intensities at each point in the chromatograms were used to calculate the absolute molecular mass from the intercept of the Debye plot, using Zimm’s model as implemented in ASTRA software (Wyatt). Fractions were analysed by gel electrophoresis and staining with SYPRO Ruby (Lonza, Cambridge, UK) or Coomassie Instant Blue according to the manufacturer’s instructions.

### Image analysis and statistics

Kymographs were generated and analysed manually using FIJI. Typically, three independent chambers were imaged using protein complexes from at least two independent assembly reactions for each experimental condition. The positions of microtubules were determined by the fluorescent tubulin signal or a projection of RNA/protein signals over the course of the movie. From each of these chambers, 5-10 microtubules were typically selected for analysis with preference given to those that were longer and better isolated from adjacent microtubules. No power analysis was used to determine sample size. Instead the sample size was chosen to allow the identification of a range of effect sizes. To avoid the risk of subconscious bias, microtubules were selected before visualising the motile properties of complexes on them. Interactions of fluorescently labelled proteins and RNA with microtubules were scored as binding events if they were ≥ 1.5s (3 frames) in duration and as processive events if they achieved predominantly minus end displacement > 500 nm (5 pixels) without significant diffusive behaviour. These parameters were chosen in advance of image acquisition following discussion with the team and all particles that fulfilled the criteria were analysed. As described previously (Schlager et al, 2014), some motile complexes changed velocity during a run. We therefore calculated velocities of individual constant-velocity segments. Run lengths were calculated from the total displacement of individual particles regardless of changes in velocity or pauses. For both velocity and run length calculations, only particles for which the entire run was observed or those with runs beginning beyond 5 µm from the microtubule minus-end were considered. When velocities and run lengths were calculated in the presence of RNA, only those complexes clearly associated with RNA were analysed. Although plots of 1 - cumulative frequency for run lengths were fitted to a one-phase exponential decay for visualisation purposes, statistical comparison of run lengths were performed on unfitted data. For Figures 1D, 2C, and 5B, ‘background’ RNA binding was quantified by generating kymographs from random microtubule-free regions of the cover slip of lengths equal to the median microtubule length of those used for analysis. For illustrative purposes, the movie and kymographs in the figures had background subtracted in Fiji with a rolling ball radius of 50 pixels. All quantitative analysis was performed on the raw data.

Statistical analyses, curve fitting, and data plotting were performed using GraphPad Prism 7.0b. A two-tailed Student’s t-test or a two-tailed Welch’s t-test was used when comparing two datasets where a Gaussian data distribution was expected, with the latter test employed in cases of unequal variance. A Mann-Whitney test was used to compare two datasets with non-Gaussian data distributions. An ANOVA test with Dunnett’s correction was used for multiple comparisons.

## AUTHOR CONTRIBUTIONS

MAM performed and analysed single molecule experiments; CD established the co-expression system for Egl and BicD proteins; CD and HTH made important contributions to establishing the *in vitro* motility assay; RM generated the Egl^dlc2pt^/BICD2 complex and performed initial experiments with it; MAM and CMJ performed SEC-MALS; MAM and SHM performed SE-AUC; MAM and SLB designed experiments and interpreted results; MAM and SLB wrote the paper; SLB supervised the project.

## ACKNOWLEDGEMENTS

We are very grateful to members of the Bullock lab, Carter lab and microscopy facility at MRC-LMB for advice and support, Andrew Carter for comments on the manuscript, Thomas Sladewski and Kathleen Trybus for sharing unpublished results, Vladimir Gelfand for the p150 antibody, and Fillip Port for the preprint template. This work was supported by the UK Medical Research Council (file reference number MC_U105178790).

## REFERENCES

Amrute-Nayak M, Bullock SL (2012) Single-molecule assays reveal that RNA localization signals regulate dynein-dynactin copy number on individual transcript cargoes. Nature Cell Biol 14: 416–423. doi:10.1038/ncb2446

Belyy V, Schlager MA, Foster H, Reimer AE, Carter AP, Yildiz A (2016) The mammalian dynein-dynactin complex is a strong opponent to kinesin in a tug-of-war competition. Nat Cell Biol 18: 1018–1024. doi:10.1038/ncb3393

Benison G, Karplus PA, Barbar E (2007) Structure and dynamics of LC8 complexes with KXTQT-motif peptides: swallow and dynein intermediate chain compete for a common site. J Mol Biol 371: 457–468. doi:10.1016/j.jmb.2007.05.046

Bieling P, Telley IA, Hentrich C, Piehler J, Surrey T (2010) Fluorescence microscopy assays on chemically functionalized surfaces for quantitative imaging of microtubule, motor, and +TIP dynamics. Methods Cell Biol 95: 555–580. doi:10.1016/S0091-679X(10)95028-0

Brautigam CA (2015) Calculations and publication-quality illustrations for analytical ultracentrifugation data. Methods Enzymol 562: 109–133. doi:10.1016/bs.mie.2015.05.001

Bullock SL, Ish-Horowicz D (2001) Conserved signals and machinery for RNA transport in *Drosophila* oogenesis and embryogenesis. Nature 414: 611–616. doi:10.1038/414611a

Bullock SL, Ringel I, Ish-Horowicz D, Lukavsky PJ (2010) A’- form RNA helices are required for cytoplasmic mRNA transport in *Drosophila*. Nat Struct Mol Biol 17: 703–709. doi:10.1038/nsmb.1813

Bullock SL, Zicha D, Ish-Horowicz D (2003) The *Drosophila hairy* RNA localization signal modulates the kinetics of cytoplasmic mRNA transport. EMBO J 22: 2484–2494. doi:10.1093/emboj/cdg230

Buxbaum AR, Haimovich G, Singer RH (2015) In the right place at the right time: visualizing and understanding mRNA localization. Nat Rev Mol Cell Biol 16: 95–109. doi:10.1038/nrm3918

Chowdhury S, Ketcham SA, Schroer TA, Lander GC (2015) Structural organization of the dynein-dynactin complex bound to microtubules. Nat Struct Mol Biol 22: 345–347. doi:10.1038/nsmb.2996

Dharan A, Opp S, Abdel-Rahim O, Keceli SK, Imam S, Diaz-Griffero F, Campbell EM (2017) Bicaudal D2 facilitates the cytoplasmic trafficking and nuclear import of HIV-1 genomes during infection. Proc Natl Acad Sci USA 114: E10707–E10716. doi:10.1073/pnas.1712033114

Dienstbier M, Boehl F, Li X, Bullock SL (2009) Egalitarian is a selective RNA-binding protein linking mRNA localization signals to the dynein motor. Genes Dev 23: 1546–1558. doi:10.1101/gad.531009

Dix CI, Soundararajan HC, Dzhindzhev NS, Begum F, Suter B, Ohkura H, Stephens E, Bullock SL (2013) Lissencephaly-1 promotes the recruitment of dynein and dynactin to transported mRNAs. J Cell Biol 202: 479–494. doi:10.1083/jcb.201211052

Edelstein A, Amodaj N, Hoover K, Vale R, Stuurman N (2010) Computer control of microscopes using microManager. Curr Prot Mol Biol. Chapter 14: Unit14 20. doi:10.1002/0471142727.mb1420s92

Grotjahn DA, Chowdhury S, Xu Y, McKenney RJ, Schroer TA, Lander GC (2018) Cryo-electron tomography reveals that dynactin recruits a team of dyneins for processive motility. Nat Struct Mol Biol. doi:10.1038/s41594-018-0027-7

Hain D, Langlands A, Sonnenberg HC, Bailey C, Bullock SL, Muller HA (2014) The *Drosophila* MAST kinase Drop out is required to initiate membrane compartmentalisation during cellularisation and regulates dynein-based transport. Development 141: 2119–2130. doi:10.1242/dev.104711

Heym RG, Zimmermann D, Edelmann FT, Israel L, Okten Z, Kovar DR, Niessing D (2013) *In vitro* reconstitution of an mRNA-transport complex reveals mechanisms of assembly and motor activation. J Cell Biol 203: 971–984. doi:10.1083/jcb.201302095

Holt CE, Schuman EM (2013) The central dogma decentralized: new perspectives on RNA function and local translation in neurons. Neuron 80: 648–657. doi:10.1016/j.neuron.2013.10.036

Hoogenraad CC, Akhmanova A (2016) Bicaudal D family of motor adaptors: linking dynein motility to cargo binding. Trends Cell Biol 26: 327–340. doi:10.1016/j.tcb.2016.01.001

Hoogenraad CC, Akhmanova A, Howell SA, Dortland BR, De Zeeuw CI, Willemsen R, Visser P, Grosveld F, Galjart N (2001) Mammalian Golgi-associated Bicaudal-D2 functions in the dynein-dynactin pathway by interacting with these complexes. EMBO J 20: 4041–4054. doi:10.1093/emboj/20.15.4041

Hoogenraad CC, Wulf P, Schiefermeier N, Stepanova T, Galjart N, Small JV, Grosveld F, de Zeeuw CI, Akhmanova A (2003) Bicaudal D induces selective dynein-mediated microtubule minus end-directed transport. EMBO J 22: 6004–6015. doi:10.1093/emboj/cdg592

Hutagalung AH, Novick PJ (2011) Role of Rab GTPases in membrane traffic and cell physiology. Phys Rev 91: 119–149. doi:10.1152/physrev.00059.2009

Huynh W, Vale RD (2017) Disease-associated mutations in human BICD2 hyperactivate motility of dynein-dynactin. J Cell Biol. doi:10.1083/jcb.201703201

Indran SV, Ballestas ME, Britt WJ (2010) Bicaudal D1- dependent trafficking of human cytomegalovirus tegument protein pp150 in virus-infected cells. J Virol 84: 3162–3177. doi:10.1128/JVI.01776-09

Kim H, Ling SC, Rogers GC, Kural C, Selvin PR, Rogers SL, Gelfand VI (2007) Microtubule binding by dynactin is required for microtubule organization but not cargo transport. J Cell Biol 176: 641–651. doi:10.1083/jcb.200608128

Liu Y, Salter HK, Holding AN, Johnson CM, Stephens E, Lukavsky PJ, Walshaw J, Bullock SL (2013) Bicaudal-D uses a parallel, homodimeric coiled coil with heterotypic registry to coordinate recruitment of cargos to dynein. Genes Dev 27: 1233–1246. doi:10.1101/gad.212381.112

Martin KC, Ephrussi A (2009) mRNA localization: gene expression in the spatial dimension. Cell 136: 719–730. doi:10.1016/j.cell.2009.01.044

Matanis T, Akhmanova A, Wulf P, Del Nery E, Weide T, Stepanova T, Galjart N, Grosveld F, Goud B, De Zeeuw CI, Barnekow A, Hoogenraad CC (2002) Bicaudal-D regulates COPI-independent Golgi-ER transport by recruiting the dynein-dynactin motor complex. Nat Cell Biol 4: 986–992. doi:10.1038/ncb891

McKenney RJ, Huynh W, Tanenbaum ME, Bhabha G, Vale RD (2014) Activation of cytoplasmic dynein motility by dynactin-cargo adapter complexes. Science 345: 337–341. doi:10.1126/science.1254198

Mofatteh M, Bullock SL (2017) SnapShot: Subcellular mRNA localization. Cell 169: 178–178 e171. doi:10.1016/j.cell.2017.03.004

Navarro C, Puthalakath H, Adams JM, Strasser A, Lehmann R (2004) Egalitarian binds dynein light chain to establish oocyte polarity and maintain oocyte fate. Nat Cell Biol 6: 427–435. doi:10.1038/ncb1122

Rapali P, Szenes A, Radnai L, Bakos A, Pal G, Nyitray L (2011) DYNLL/LC8: a light chain subunit of the dynein motor complex and beyond. FEBS J 278: 2980–2996. doi:10.1111/j.1742-4658.2011.08254.x

Reck-Peterson SL, Yildiz A, Carter AP, Gennerich A, Zhang N, Vale RD (2006) Single-molecule analysis of dynein processivity and stepping behavior. Cell 126: 335–348. doi:10.1016/j.cell.2006.05.046

Redwine WB, DeSantis ME, Hollyer I, Htet ZM, Tran PT, Swanson SK, Florens L, Washburn MP, Reck-Peterson SL (2017) The human cytoplasmic dynein interactome reveals novel activators of motility. eLife 6. doi:10.7554/eLife.28257

Reichhart JM, Ferrandon D (1998) Green balancers. Drosoph Inf Serv 81: 201–202.

Schlager MA, Hoang HT, Urnavicius L, Bullock SL, Carter AP (2014) *In vitro* reconstitution of a highly processive recombinant human dynein complex. EMBO J 33: 1855–1868. doi:10.15252/embj.201488792

Schuck P (2003) On the analysis of protein self-association by sedimentation velocity analytical ultracentrifugation. Analytical Biochem 320: 104–124. doi:

Sharp DJ, Brown HM, Kwon M, Rogers GC, Holland G, Scholey JM (2000) Functional coordination of three mitotic motors in *Drosophila* embryos. Mol Biol Cell 11: 241–253. doi:

Short B, Preisinger C, Schaletzky J, Kopajtich R, Barr FA (2002) The Rab6 GTPase regulates recruitment of the dynactin complex to Golgi membranes. Curr Biol 12: 1792–1795. doi:

Sladewski TE, Bookwalter CS, Hong MS, Trybus KM (2013) Single-molecule reconstitution of mRNA transport by a class V myosin. Nat Struct Mol Biol 20: 952–957. doi:10.1038/nsmb.2614

Sladewski TE, Billington N, Ali M, Bookwater CS, Lu H, Krementsova EB, Schroer TA and Trybus KM (2018) Recruitment of two dyneins to an mRNA-dependent Bicaudal D transport complex. bioRxiv, doi.org/10.1101/273755

Soundararajan HC, Bullock SL (2014) The influence of dynein processivity control, MAPs, and microtubule ends on directional movement of a localising mRNA. eLife 3: e01596. doi:10.7554/eLife.01596

Splinter D, Razafsky DS, Schlager MA, Serra-Marques A, Grigoriev I, Demmers J, Keijzer N, Jiang K, Poser I, Hyman AA, Hoogenraad CC, King SJ, Akhmanova A (2012) BICD2, dynactin, and LIS1 cooperate in regulating dynein recruitment to cellular structures. Mol Biol Cell 23: 4226–4241. doi:10.1091/mbc.E12-03-0210

Stuurman N, Haner M, Sasse B, Hubner W, Suter B, Aebi U (1999) Interactions between coiled-coil proteins: *Drosophila* lamin Dm0 binds to the bicaudal-D protein. Eur J Cell Biol 78: 278–287. doi:

Terawaki S, Yoshikane A, Higuchi Y, Wakamatsu K (2015) Structural basis for cargo binding and autoinhibition of Bicaudal-D1 by a parallel coiled-coil with homotypic registry. Biochem Biophys Res Comm 460: 451–456. doi:10.1016/j.bbrc.2015.03.054

Urnavicius L, Lau CK, Elshenawy MM, Morales-Rios E, Motz C, Yildiz A, Carter AP (2018) Cryo-EM shows how dynactin recruits two dyneins for faster movement. Nature 554: 202–206. doi:10.1038/nature25462

Urnavicius L, Zhang K, Diamant AG, Motz C, Schlager MA, Yu M, Patel NA, Robinson CV, Carter AP (2015) The structure of the dynactin complex and its interaction with dynein. Science 347: 1441–1446. doi:10.1126/science.aaa4080

Van De Bor V, Hartswood E, Jones C, Finnegan D, Davis I (2005) *gurken* and the I factor retrotransposon RNAs share common localization signals and machinery. Dev Cell 9: 51–62. doi:10.1016/j.devcel.2005.04.012

Wilkie GS, Davis I (2001) *Drosophila wingless* and pair-rule transcripts localize apically by dynein-mediated transport of RNA particles. Cell 105: 209–219. doi:

Zhang K, Foster HE, Rondelet A, Lacey SE, Bahi-Buisson N, Bird AW, Carter AP (2017) Cryo-EM reveals how human cytoplasmic dynein is auto-inhibited and activated. Cell 169: 1303–1314 e1318. doi:10.1016/j.cell.2017.05.025

